# Genotoxic antibody-drug conjugates combined with Bcl-xL inhibitors enhance therapeutic efficacy in metastatic castration-resistant prostate cancer

**DOI:** 10.1101/2025.09.05.674562

**Authors:** Galina Semenova, Sander Frank, Ruth Dumpit, Wanting Han, Ilsa Coleman, Roman Gulati, Colm Morrissey, Michael C. Haffner, Peter S. Nelson, John K. Lee

**Author notes:** Corresponding authors: Galina Semenova, UCLA Jonsson Comprehensive Cancer Center, 700 Tiverton Drive, Factor Building 8-240, Los Angeles, CA 90095.; Peter S. Nelson, Fred Hutchinson Cancer Center, Seattle, WA 98109.; John K. Lee, UCLA Jonsson Comprehensive Cancer Center, 700 Tiverton Drive, Factor Building 8-240, Los Angeles, CA 90095.

## Abstract

Metastatic castration-resistant prostate cancer (mCRPC) is an aggressive subtype of prostate cancer (PC) without curative treatments. Antibody-drug conjugates (ADCs) emerged as promising cancer therapeutics that selectively deliver cytotoxic agents (payloads) to the tumors. Although ADCs have been successfully applied in the treatment of hematological and solid tumors, ADC monotherapy has not demonstrated durable responses in mCRPC and the mechanisms of PC resistance to ADCs have not been thoroughly investigated. Our study aimed to improve ADC efficacy using a new integrated approach for custom ADC design and multiplexing. To nominate rational combinations of ADC targets and ADC payloads, we (1) examined protein co-expression of three clinically relevant surface antigens— B7 homolog 3 (B7-H3), prostate specific membrane antigen (PSMA), and six-transmembrane epithelial antigen of prostate-1 (STEAP1)—in a series of human mCRPC samples and (2) screened established ADC payloads and their combinations in mCRPC cell lines with different phenotypes. We identified synergistic interactions between DNA-damaging payloads and Bcl-xL inhibitor A-1331852 as well as their coordinated induction of the intrinsic apoptosis pathway. The functional relevance of isolated p53 loss and impaired PC responses to three genotoxic ADCs (B7-H3-seco-DUBA, PSMA-SG3249, and STEAP1-DXd) and their combinations with A-1331852 was established using genetic knockout models. Lastly, we found enhanced *in vivo* antitumor activity in mCRPC by combining the clinically relevant agents B7-H3-seco-DUBA (vobramitamab duocarmazine) and A-1331852. Collectively, our findings provide rationale for the development of ADC therapies combining genotoxic payloads with Bcl-xL inhibitors for mCRPC.

**Significance:** B7-H3, PSMA, and STEAP1 targeted ADC therapies combining genotoxic payloads with Bcl-xL inhibitors induce p53-dependant apoptotic cell death in mCRPC, providing a clinically viable strategy for the treatment of advanced prostate cancer.

## INTRODUCTION

Prostate cancer (PC) is the most frequently diagnosed non-cutaneous malignancy and is the second leading cause of cancer death in males (1). Localized PC is curable by surgery or radiation therapy; however, metastatic PC remains incurable. While these tumors initially respond to androgen deprivation therapy, nearly all will progress to lethal, metastatic castration-resistant PC (mCRPC). Effective application of systemic chemotherapy to eradicate mCRPC is restricted by dose-limiting toxicities and the inevitable development of resistance (2,3), suggesting the need for improved drug delivery to the tumor site and diversification of applied cytotoxic agents.

Antibody-drug conjugates (ADCs) are a new generation of targeted therapeutics that combine the high specificity of monoclonal antibodies with the potency of chemotherapeutic drugs (4). An ADC is comprised of three parts: 1) an antibody that binds to an antigen on the target cell surface, 2) a cytotoxic small molecule (payload), and 3) a linker that enables conjugation of the payload to the antibody and plays a critical role in the release of the payload upon delivery to the tumor site (5). These weaponized antibodies can be tailored for specific cancer types, and many ADCs have demonstrated activity against treatment-refractory cancers (4).

Despite the successes of ADCs in the treatment of breast and urothelial tumors, the clinical application of ADCs to mCRPC has thus far been unsuccessful (6). Limited responses to ADC monotherapy in mCRPC may be explained by tumor antigen heterogeneity, on-target and off-target systemic toxicity, and by the development of tumor resistance to ADC components (6,7). Under continuous treatment pressure, cancer cells can acquire resistance to both antibody targeting by down-regulation/mutation of cell surface antigens and to the ADC payload through, for instance, the upregulation of drug-efflux transporters (7). Therefore, the rational selection of ADC targets and linker-payloads as well as the development of synergistic ADC combinations have the potential to enhance the therapeutic window for the effective treatment of mCRPC.

To date, several preclinical and clinical studies have assessed the efficacy of ADCs combined with other types of anticancer drugs, including endocrine therapy, chemotherapy, targeted molecular agents, and immunotherapy for the treatment of solid tumors (8). These combination therapies have demonstrated some success in improving antitumor efficacy; however, research on PC combination therapies involving ADCs remains limited.

In the present study, we evaluated the benefits of ADC combinations that leverage synergism between two distinct and complementary ADC payloads. To design ADCs for combination mCRPC therapy with improved efficacy and specificity, we first analyzed the distribution of clinically relevant PC targets: prostate specific membrane antigen (PSMA, *FOLH1*) (9,10), six-transmembrane epithelial antigen of prostate-1 (STEAP1) (11,12), and B7 homolog 3 (B7-H3, *CD276*) (13–15) in mCRPC and normal tissues and proposed to use these antigens in pairs for dual ADC targeting. Next, we screened established ADC payloads in pairwise combinations and identified synergistic interactions between DNA-damaging payloads and the Bcl-xL inhibitor A-1331852. Finally, we confirmed the activity of synergistic payloads administered through combination ADC therapy using *in vitro* and *in vivo* PC models. Collectively, our findings indicate that combining ADCs that enable simultaneous delivery of two synergistic payloads is a promising approach to potentiate therapeutic activity in mCRPC.

## RESULTS

### B7-H3(CD276), PSMA(FOLH1) and STEAP1 are co-expressed in metastatic prostate cancers

We sought to enhance the anti-tumor effects of prostate cancer therapeutics by directing the cytotoxic payloads via antibody delivery and using combination strategies that exploit multiple target antigens and synergistic payloads. Prior findings show that co-administration of two ADCs targeting the same antigen may lead to binding competition and reduced efficiency of payload delivery (16). Thus, we proposed to use two different antigens to simultaneously co-target tumor cells with ADCs for the treatment of mCRPC. Such target pairs should be expressed on the surface of the same PC cells though have limited normal tissues co-expression.

Several surface antigens have been shown to be highly expressed by prostate cancers and have been previously employed as ADC targets for mCRPC. The most well-established antigens include PSMA (17–19), STEAP1 (11,12,20) and B7-H3 (15,21,22); however, to exploit combinatorial effects, it is essential to confirm that these antigens are expressed by the same cells in mCRPC. To address this issue, we conducted multiplexed immunofluorescent (mIF) staining for B7-H3, PSMA, and STEAP1 in a clinically and histologically annotated case series of mCRPC tumors (**Figure 1A, Table S3**). Tumor tissues were collected at rapid autopsy through the University of Washington Tissue Acquisition Necropsy (UW TAN) program (23). Tumor sections were organized within a tissue microarray (TMA) comprised of 181 tumors from 58 patients. A total of five tumors were not analyzed due to insufficient tumor content, leaving 176 evaluable tumors (514 individual cores) from 58 patients. Tumors were classified into four phenotypic subtypes based on immunohistochemical staining for androgen receptor (AR) signaling and neuroendocrine (NE) markers as previously described (24,25): AR-active prostate cancer (ARPC or AR+/NE-), amphicrine carcinoma (AR+/NE+), double negative mCRPC (AR-/NE-), neuroendocrine prostate cancer (NEPC or AR-/NE+). Integrated H-scores combining the frequency and intensity of staining were used to evaluate single antigen staining (**Figure 1**). For dual antigen staining analysis, quantification of the percentage of double-positive cells was applied (**Figure 2**).

**Figure 1.**
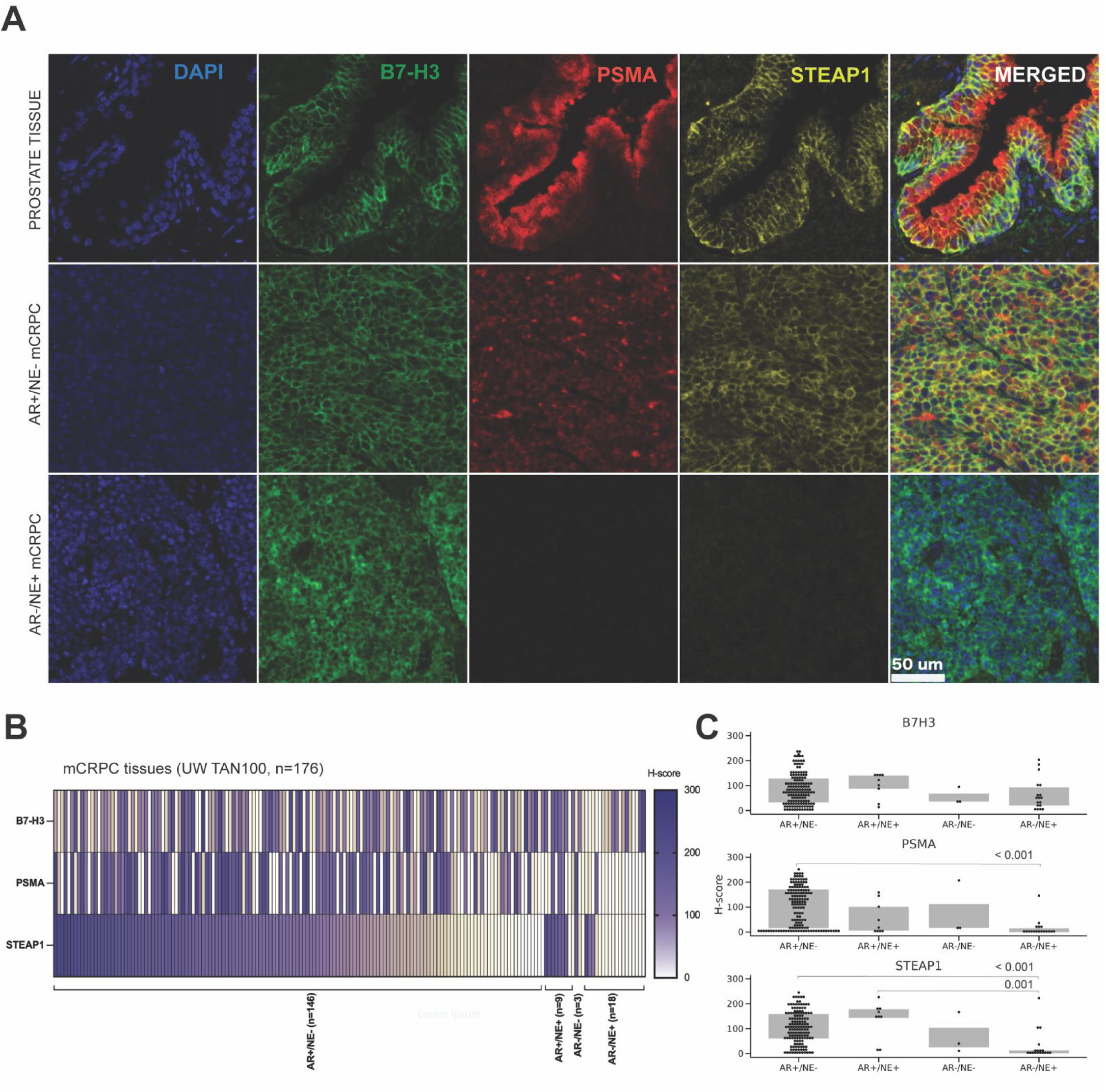
B7-H3, PSMA, and STEAP1 are expressed in the majority of AR regulated mCRPCs. (A) Representative TMA images of human prostate tissue (FDA999 L206) and mCRPC tissues (UW TAN100) with membranous B7-H3, PSMA, STEAP1, and nuclear DAPI staining. (B) Heatmap of individual H-scores of 176 mCRPC tissues. (C) Averaged H-scores of AR+/NE-, AR+/NE+, AR+/NE-, and NEPC tissue samples. Wilcoxon-Mann-Whitney rank sum test p values are shown for plot C.

**Figure 2.**
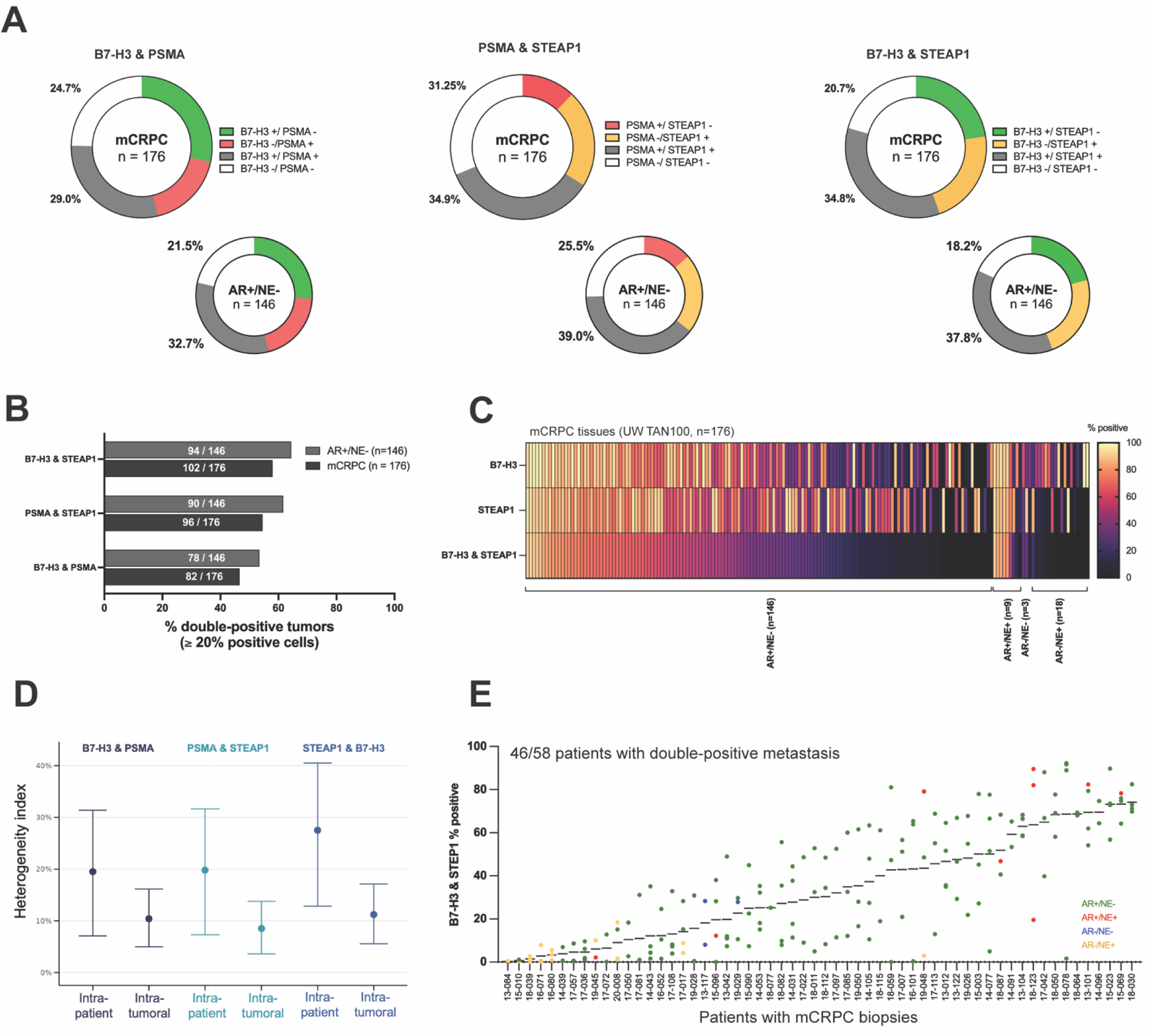
mCRPC cells co-express ADC target pairs PSMA and STEAP1/ B7-H3 and PSMA/ B7H3 and STEAP1. (A) Averaged percentage double-positive and double-negative cells in all mCRPC and specifically in AR+/NE-tumors. (B) The percentage of mCRPC or AR+/NE-tumors co-staining for each antigen pair. Heatmap showing percents of cells staining positively for B7-H3, STEAP1, or both in individual mCRPC tumors (columns, n=176). (D) Hypergeometric mean (95% CI) of antigen co-expression heterogeneity indices across different metastatic sites in a given patient (intra-patient heterogeneity) and within a metastatic site (intra-tumoral heterogeneity). (E) Distribution of B7-H3 and STEAP1 double-positive cells in 176 metastatic tumors within and between 58 patients from UW TAN cohort. Each dot represents a tumor sample; the color codes indicate the molecular subtype – AR+/NE-(green), AR+/NE+ (red), AR-/NE-(blue), and AR-/NE+ (yellow).

We observed B7-H3, PSMA, and STEAP1 staining heterogeneity across the four mCRPC phenotypic groups (**Figure 1B**). Consistent with prior reports (12,17), STEAP1 and PSMA H-scores were significantly greater in AR+/NE-tumors compared to NEPC; however, B7-H3 reactivity was observed in both AR-active and NE tumors (**Figure 1C**). These findings are consistent with the corresponding B7-H3/*CD276*, PSMA/*FOLH1*, and *STEAP1* gene expression analysis in three cohorts determined by RNAseq: 270 Stand Up To Cancer (SU2C) international dream team mCRPC biopsies, 172 mCRPC samples from the UW TAN program, and 126 LuCaP PC patient-derived xenograft (PDX) tumors (**Figure S1**).

B7-H3, PSMA and STEAP1 IF staining was evaluated pairwise at single-cell resolution, both within tumors and across tumors for individual patients. To assess the fraction of cells co-expressing two antigens, the percentage of double-positive cells was calculated in all tumor specimens (**Figure 2A**). We determined that approximately one-third of all mCRPC cells stained positively for the combination of PSMA and STEAP1, B7-H3 and PSMA, or B7-H3 and STEAP1. Within the AR+/NE-mCRPC cohort 32.7-39% of cells were double-positive. The largest antigen overlap was observed between PSMA and STEAP1 where 39% of cells comprising AR+/NE-tumors stain for both, while the greatest antigen coverage was achieved with the B7-H3 and STEAP1 pair, with only 18.2% cells in AR+/NE-tumors staining for neither antigen (81.8% cells stain for either B7-H3 or STEAP1). Overall, 20% of mCRPC cells across all phenotype groups, and 25% of AR+/NE-cells stained positively for all three antigens.

We next quantified the frequency at which tumors contained a large fraction of double-positive cells. Tumor cores with ≥20% cells co-staining for any antigen pair were considered double-positive. About half of the tumors were double-positive for at least one antigen pair, and most of the tumors were B7-H3 and STEAP1 positive: 58% (102/176) mCRPC and 64.4% (94/146) AR+/NE-tumors (Figure 2B-C, S2A-B). B7-H3 and STEAP1 pair also demonstrated the highest estimated staining heterogeneity (intra-patient 27.4% and intra-tumoral 11.2%), followed by B7-H3 and PSMA (19.5% and 9.4%) and PSMA and STEAP1 (19.4% and 8.4%) (**Figure 2D**). 46/58 (79.3%) mCRPC patients had at least one B7-H3 and STEAP1 positive tumor, 39/58 (67.2%) patients demonstrated at least one PSMA and STEAP1 positive tumor, and 34/58 (58.6%) patients harbored at least one B7-H3 and STEAP1 positive tumor (**Figure 2E, S2B**). All tumors stained positive for STEAP1 and B7-H3 in 17 patients, for PSMA and STEAP1 in 21 patients, and for B7-H3 and PSMA in 15 patients. Collectively, these assessments of inter-tumor and inter-individual homogeneity and heterogeneity provide data to inform combinatorial strategies designed to target the same or divergent cell populations within a patient and how to deploy targeted agents with systemic delivery.

Antigen target co-expression was also evaluated in a human benign tissue TMA (**Figure 1A, S3**). We found that luminal cells of the prostate and mammary gland stained positively for all three antigens: STEAP1, B7-H3, and PSMA. Kidney tubular cells and small intestine enterocytes co-stained for STEAP1 and PSMA, while urothelial cells and colon enterocytes showed low intensity B7-H3 and STEAP1 staining.

Overall, these data collectively show that either of three antigen pairs demonstrate high levels of double-positivity in mCRPC tumors, particularly in the AR+/NE-phenotype which is the most common subtype of CRPC, and very low levels of double-positivity in normal tissues, suggesting that these antigen pairs may be compelling targets for combination ADC therapy.

### Nomination of synergistic interactions between established ADC payloads in PC

To date, the majority of ADCs are comprised of payloads with similar mechanisms of action (MoA) that fall into two major classes: microtubule targeting or DNA damaging agents. This raises concerns about the development of cross-resistance with systemic standard-of-care treatments or with other ADCs if used sequentially. We therefore asked if rational selection of the payload MoA for a specific molecular subtype of PC could improve therapeutic response and mitigate ADC resistance.

Recent efforts have yielded ADC payloads using different MoA beyond conventional agents targeting DNA or microtubules (26). The development of novel payloads provides opportunities for ADC co-targeting, as combinations of agents with different MoA are predicted to have the highest probability of overcoming resistance without overlapping toxicities. Payload combinations that offer additive or synergistic cytotoxic effects on cancer cells can potentially increase ADC efficacy and reduce effective therapeutic doses.

To evaluate the PC cell response to individual ADC payloads and to nominate synergistic payload interactors, we designed a drug screen with 23 small molecules representing three major classes of ADC payloads: microtubule disrupting drugs (MDDs), DNA-damaging drugs (DDDs), and innovative drugs (IDs) targeting RNA synthesis (thailanstatins, amatoxins), NAD synthesis (NAMPT inhibitors), or apoptosis (Bcl-xL inhibitor). Each payload class included 4-5 pharmacological groups with different MoA and 1-3 analogous members per group (**Figure 3A**). The 23 agents were evaluated at two working concentrations (high and low) which were individually selected for each compound based on effective concentrations reported prior (**Figure 3A, Table S1**). We also evaluated 13 out of 23 compounds (one representative from each payload group) in pairs at low working concentrations, resulting in 78 total pairwise drug combinations (**Figure 3A**). Twenty-three payloads and active payload derivatives were tested in four mCRPC cell lines representing different phenotypic subtypes: C4-2B (AR+/NE-), 22Rv1 (AR+/NE+), LuCaP 176 (AR-/NE-), and MSKCC EF1 (AR-/NE+) (**Figure 3A-B, S4, Table S4**). Agents from all three payload classes effectively killed AR-regulated cells C4-2B and 22Rv1 at high and low doses (**Figure 3B, S4**). LuCaP 176 cell line demonstrated weaker responses to many drugs and in particular to DDDs compared to other lines. The NEPC cell line MSKCC EF1 showed moderate responses to MDDs and DDDs; however, it was sensitive to several IDs including the Bcl-xL inhibitor A-1331852, suggesting potential utility of this payload in the development of ADCs for NEPC. When administered at high doses, cytotoxic effects of DDDs were significantly greater than the effects of MDDs across all mCRPC models (**Figure S4**).

**Figure 3.**
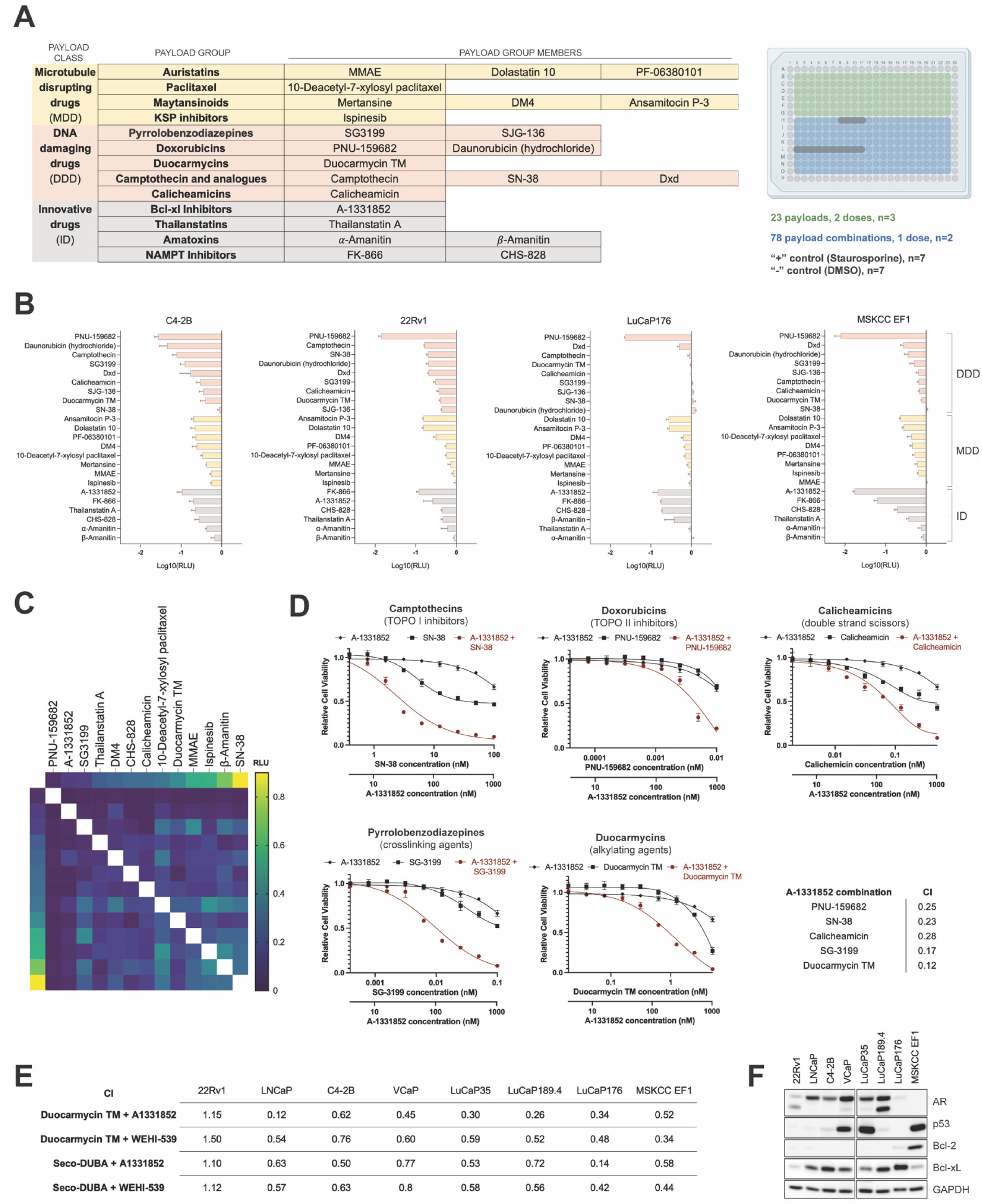
Screening of payload combinations identifies synergistic interactors. (A) The list of cytotoxic agents tried in the drug screen (left) and the layout of the 384-well screen master plate (right) containing 23 individual drugs, 78 drug combinations, and the positive and negative (vehicle) controls. (B) Normalized responses of C4-2B, 22Rv1, LuCaP176, and MSKCC EF1 to individual payloads exposure (low dose). Compounds are organized by the payload class – DNA damaging drugs, DDD (salmon), microtubule disrupting drugs, MDD (yellow), and innovative drugs, ID (gray). Results are expressed as log 10 relative luminescence units (RLU). (C) Heatmap demonstrating normalized responses of C4-2B cells to single payloads (data points on the left and upper edge) and payload pairs (data points in the center of the heatmap). (D) Dose-response to Bcl-xL inhibitor A-1331852, DDDs SN-38, PNU-159682, Calicheamicin, SG-3199, Duocarmycin TM, and 5 combinations of DDDs with A-1331852 in LNCaP cells. (E) Combination indices (CI) scored for 4 drug combinations in 8 PC cell lines. (F) Immunoblot analysis of AR, p53, Bcl-2 and Bcl-xL protein levels in 8 PC cell lines.

We next assessed the effects of drug/payload combinations. We observed significant differences in the response to drug combinations between the four PC models (**Figure S5A**). Payload pairs for the synergy validation were selected from the interactors with different MoA that exhibited PC cell toxicity greater than the amplification of their individual effects in at least two cell lines (**Figure 3C, S5A**). Potential synergistic interactions were observed between DDDs and IDs, as well as between the members of ID group (**Figure S5B**). These pairs were prioritized and further validated using a dilution series of payloads alone and in combination. Drug additivity or synergism between payloads was analyzed in the *in vitro* cell viability assays using Chou-Talalay method (27). Interactors with combination indices (CI) < 1.0 were considered synergistic. Based on these validation studies, a highly enriched group of synergistic interactions was identified between the Bcl-xL inhibitor A-1331852 and DNA-damaging agents from all five DDD groups tested in the screen: SN-38 (TOPO I inhibitor), PNU-159682 (TOPO II inhibitor), Calicheamicin (double strand DNA scissor), SG-3199 (DNA crosslinking agent), and Duocarmycin TM (DNA alkylating agent) (**Figure 3D**).

To confirm on-target drug effects, synergism between Duocarmycin TM and A-1331852 was further verified using the drug analogs WEHI-539 (Bxl-xL inhibitor) and *seco*-DUBA (duocarmycin analog) in a panel of eight PC cell lines (**Figure 3E, S6, S7**). All but one of the 8 models tested demonstrated a CI of <1 with each combination. The 22Rv1 model was the exception and these results can likely be attributed to the low Bcl-xL protein level in this cell line (**Figure 3F**) as Bcl-xL (*BCL2L1*) expression may be necessary for the cell response to the proposed payload combination. These findings suggest that co-targeting mCRPC with two ADCs bearing DNA damaging agents and Bcl-xL inhibitors as payloads may improve tumor responses to ADC therapy.

### Induction of p53-mediated apoptosis through DNA damage and Bcl-xL inhibition in PC

The synergism between DDDs and A-1331852 can be explained by the interplay between DNA damage and Bcl-xL function in the intrinsic p53-dependent apoptosis pathway (28). In response to DNA damage p53 is rapidly activated and stabilized through posttranslational modifications, that engage several cell death programs including mitochondrial cell death(29). Pro-survival Bcl-2 family proteins, including Bcl-2 and Bcl-xL, are localized in the mitochondria and prevent apoptosis by inhibiting permeabilization of the outer mitochondrial membrane (30). Our previous report (31) and current data indicate that while Bcl-2 is overexpressed in mCRPCs with NE phenotype, Bcl-xL is abundant in both AR-active tumors and NEPCs (**Figure 3F, 4A**).

In several cancer types p53 loss-of-function promotes resistance to DNA damaging chemo- and radio-therapeutics (32). In this context, we determined that over a third of AR+/NE-tumors in the SU2C cohort demonstrate biallelic loss of *TP53* (**Figure 4A, S8**). We queried how functional *TP53* knockout (KO) would affect PC cell responses to genotoxic payloads and their combinations with the Bcl-xL inhibitor A-1331852. To compare the cytotoxicity of DDDs in isogenic cell models we performed CRISPR/Cas9 knock out (KO) of *TP53* to generate three isogenic PC models with and without functional *TP53*: LNCaP (single clone (33)), C4-2B (single clone), and LuCaP 189.4 (KO pool). Parental and *TP53* KO cells were treated with SN-38, PNU-159682, Calicheamicin, SG-3199, and Duocarmycin TM for 72 hours (**Figure 4B**). In all three isogenic cell line pairs we observed a weaker response to DDDs in *TP53* KO cells compared to the parental cells with intact *TP53*. Likewise, parental cells were more sensitive to Duocarmycin TM and A-1331852 combination (**Figure 4C**). Induction of apoptosis in p53-proficient cells after a 24 hour exposure to Duocarmycin TM and A-1331852 was confirmed by assaying PARP1 and caspase 3 cleavage (**Figure 4D**). We also observed an increase in p53 protein levels in the p53 positive cells treated with Duocarmycin TM, which is indicative of p53 stabilization following DNA damage (**Figure 4D**). These findings imply that, along with Bcl-xL (*BCL2L1*) expression, p53 proficiency might be a contributing factor to PC cell response to the combination of DDDs and Bcl-xL inhibitor payloads.

**Figure 4.**
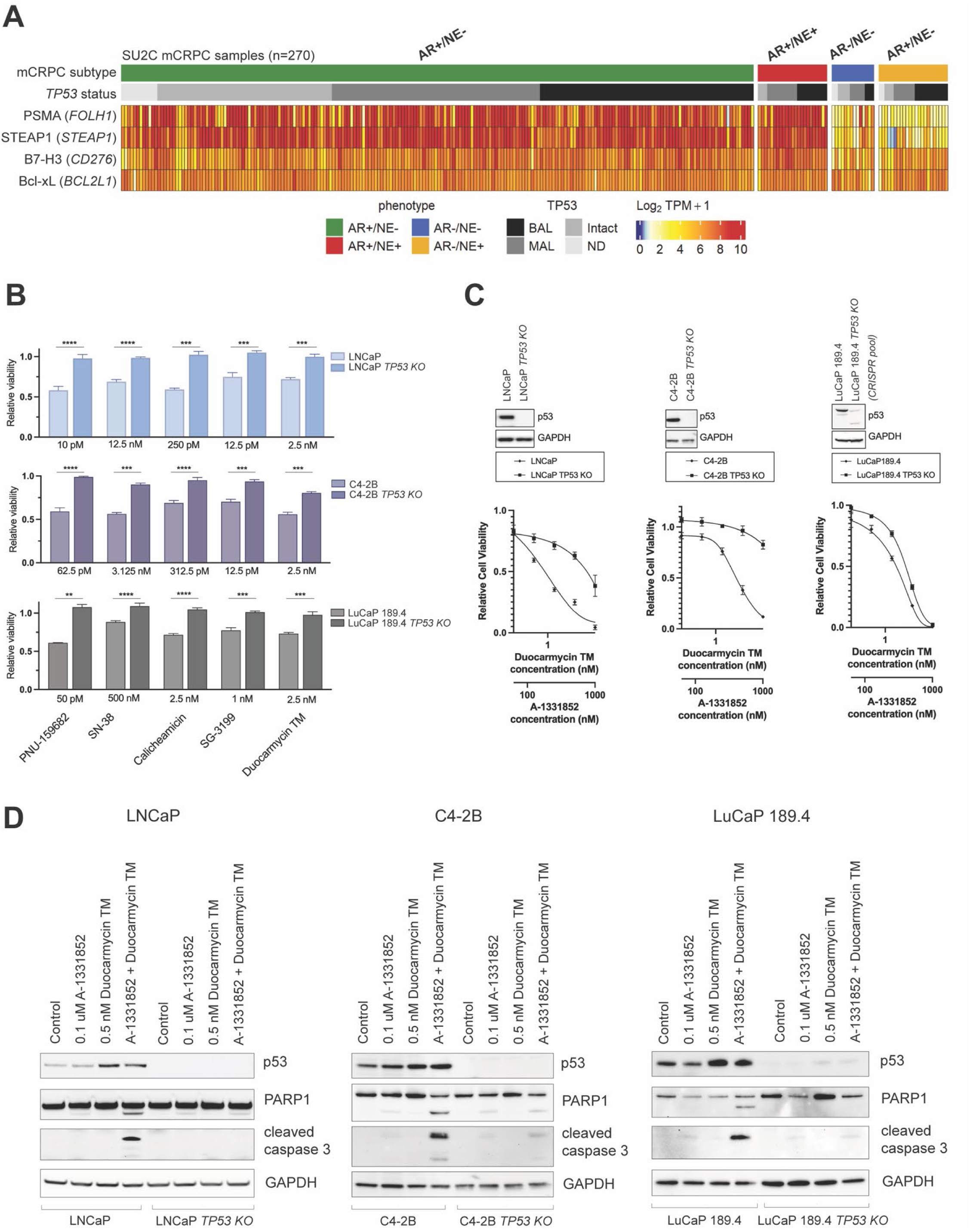
Combination of Duocarmycin TM with A-1331852 induces apoptosis in prostate cancer lines expressing p53(*TP53) and* BCL-XL*(BCL2L1)*. (A) Heatmap showing *FOLH1, STEAP1, CD276,* and *BCL2L1* transcript abundance, as well as *TP53* genomic status in 270 SU2C mCRPC specimens. Transcript levels are shown as Log_2_ TPM + 1. BAL – biallelic loss, MAL – monoallelic loss, ND – no data. (B) Relative viability of PC cells lines expressing wild type *TP53* (LNCaP, C4-2B, LuCaP189.4) and isogenic *TP53 KO* cells exposed to DNA-damaging agents for 72 hrs. Data shown as mean with standard deviation. Significance determined using unpaired t-test. (C) Immunoblot analysis of p53 protein levels and dose-response of isogenic lines to Duocarmycin TM and A-1331852 combination. Immunoblot analysis of p53, PARP1, and cleaved caspase 3 in isogenic cell lines treated with DMSO (Control), A-1331852, Duocarmycin TM, or the combination of A-1551852 and Duocarmycin TM for 24 hrs.

### Genotoxic ADCs combined with the Bcl-xL inhibitor A-1331852 enhance cytotoxicity

To verify that payload synergism can be translated into ADC combination regimens we combined ADCs bearing DNA-damaging payloads with unconjugated Bcl-xL inhibitor A-1331852. As conjugation of A-1331842 to an antibody required significant chemical synthesis, this drug was used as a free systemic agent for a proof-of-concept. Three different genotoxic ADCs targeting B7-H3, PSMA, or STEAP1 were tested singly and in combination with A-1331852 (**Table S5**). C4-2B and 22Rv1 cell lines with intact B7-H3 (*CD267*), PSMA (*FOLH1*), and STEAP1 and isogenic paired lines with gene deletions of these proteins (PSMA KO and STEAP1 KO) were engineered to confirm antibody specificity and ADC responses (**Figure 5A**).

**Figure 5.**
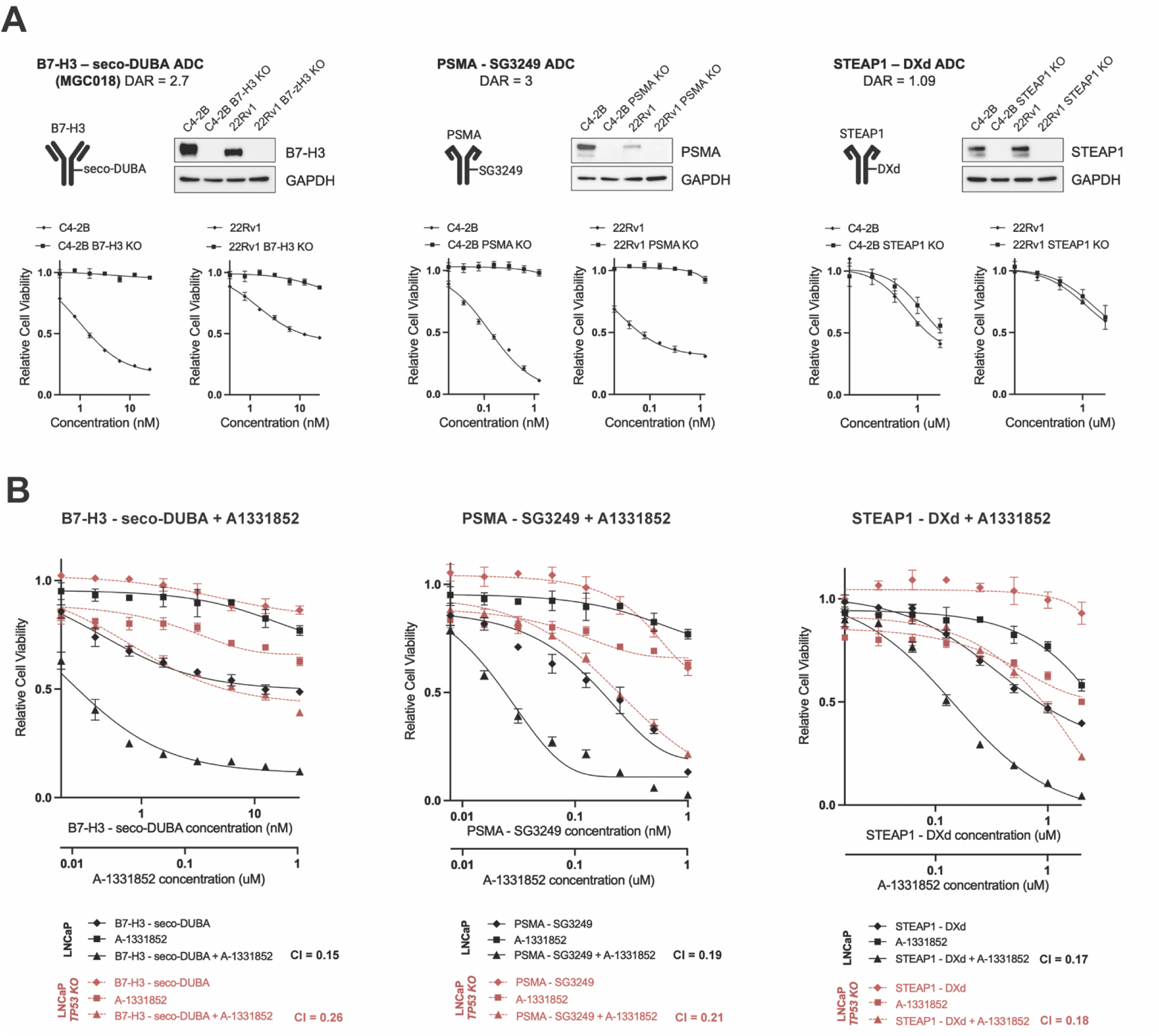
ADCs bearing genotoxic payloads synergize with A-1331852 *in vitro*. (A) Diagrams of B7-H3-seco-DUBA, PSMA -SG3249, and STEAP1-DXd ADCs. Immunoblots showing B7-H3, PSMA, and STEAP1 knockouts in isogenic C4-2B and 22Rv1 cell lines. Differential response of antigen expressing cell lines to B7-H3-seco-DUBA, PSMA-SG3249, and STEAP1-DXd ADCs (4 days exposure). (B) Synergistic interactions between genotoxic ADCs and Bcl-xL inhibitor A-1331852 in *TP53 WT* and *TP53 KO* LNCaP cells. Plots show mean −/+ standard deviation (n=3). CI – combination index.

B7-H3-seco-DUBA (MGC018, vobramitamab duocarmazine, vobra duo), a clinical grade ADC targeting B7-H3 and bearing the genotoxic seco-DUBA duocarmycin prodrug payload (21), was provided by MacroGenics. To produce additional genotoxic ADCs, we synthesized single chain antibodies (scFv-Fc) binding PSMA (J591 biosimilar) and STEAP1 (vandortuzumab biosimilar) (**Figure S9**). The binding specificity of PSMA scFv-Fc was confirmed by flow cytometry in parental and PSMA KO cells while specificity of STEAP1 scFv-Fc was confirmed in parental and STEAP1 KO cells (**Figure S9A**). No cytotoxic effects of naked PSMA scFv-Fc or STEAP1 scFv-Fc were observed in C4-2B or 22Rv1 cells (**Figure S9B**). After ensuring antigen specificity, PSMA scFv-Fc was conjugated to Tesirine (SG3249), a pyrrolobenzodiazepine dimer payload, resulting in PSMA-SG3249 ADC with a drug-to-antibody ratio (DAR) of 3. STEAP1 scFv-Fc was conjugated to the camptothecin analog Deruxtecan (DXd) resulting in STEAP1-DXd with a DAR of 1.09. Enzymatically cleavable linkers were used in all three ADC designs (**Table S5**).

B7-H3-seco-DUBA, PSMA-SG3249, and STEAP1-DXd demonstrated reasonable selective toxicity against cells expressing target antigen (**Figure 5A**) and were subsequently applied to *TP53* wildtype (WT) and *TP53* KO LNCaP cells with and without A-1331852. Synergistic interactions between genotoxic ADCs and A-1331852 was observed in both cell models and in all ADC combinations tested (**Figure 5B**); however, the responses were greater in p53-proficient cells. These findings indicate that the addition of Bcl-xL inhibitor A-1331852 to clinically relevant ADCs bearing DNA damaging payloads can significantly improve PC cell responses to these ADCs *in vitro*.

### Antitumor activity of MGC018 and A-13318562, alone or in combination, in p53-deficient and p53-proficient mCRPC

The antitumor activity of B7-H3-seco-DUBA ADC (MGC018) and its combination with unconjugated A-1331852 *in vivo* was examined using C4-2B and C4-2B *TP53 KO* cell-derived xenograft (CDX) tumors established in male NSG mice. Mice were treated with vehicle, B7-H3-targeted ADC, A-1331852, or the combination of B7-H3-targeted ADC and A-1331852 for a period of two weeks. Tumor growth was monitored for an additional two weeks after the end of treatment. MGC018 at 3 mg/kg was administered weekly via intraperitoneal injection, A-1331852 at 25 mg/kg was administered via oral gavage 5 times per week.

Overall, we observed a greater and more rapid inhibition of tumor growth in response to MGC018 in p53-proficient tumors (**Figure 6A, S10A**). The addition of Bcl-xL inhibitor significantly enhanced antitumor activity in combination with MGC018 in both C4-2B and C4-2B *TP53 KO* CDXs (**Figure 6A, S10A**). In the C4-2B experiment, tumor volume in the control group grew 9-10% per day. The Bcl-xL inhibitor A1331852 reduced the average daily growth rate to 6-7%. MGC018 reduced the average daily growth rate to 2-5%, while the combination of both drugs reduced growth to 1-2%. In the C4-2B *TP53* KO experiment, tumors were smaller, and the volume in the control group grew on average 6-7% per day. The Bcl-xL inhibitor A1331852 reduced the average daily growth rate to 5-6%. MGC018 reduced the average daily growth rate to 3-5%. The combination of both MGC018 and A-1331852 reduced the daily growth rate to 1-2%. In C4-2B CDXs adding A1331852 to MGC018 reduced the average daily growth rate by 2-3% (95% CI, p<0.001), while in C4-2B *TP53* KO model it reduced the average daily growth rate by 1-3% (95% CI, p=0.014) (**Figure S10A**).

**Figure 6.**
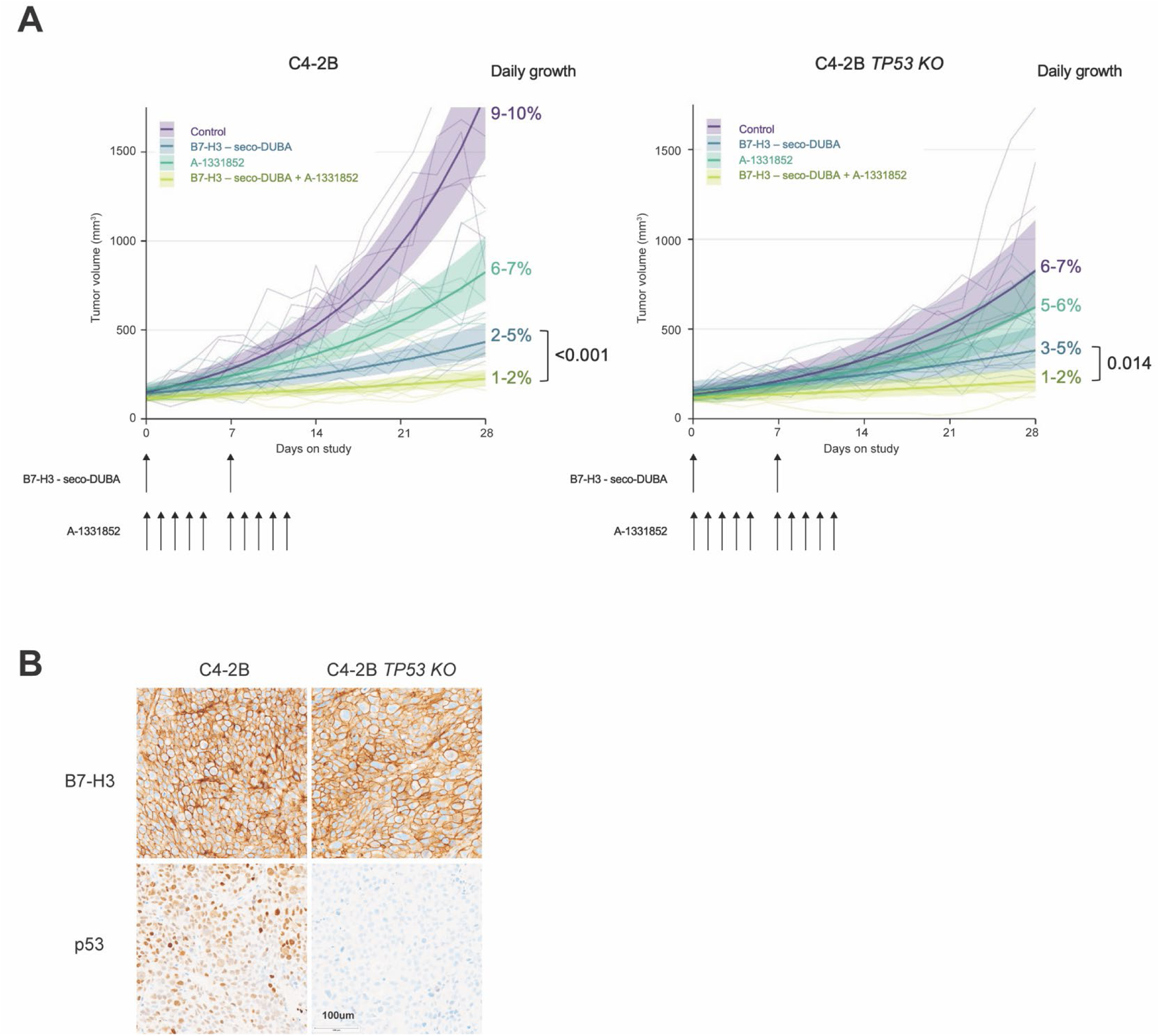
Combination B7-H3-seco-DUBA ADC with systemic A-1331852 inhibits the growth of p53 proficient and p53 deficient mCRPCs. (A) Volumetric changes of C4-2B and C4-2B *TP53 KO* xenograft tumors in NSG mice (n=6-7) exposed to vehicle alone, B7-H3-seco-DUBA ADC (3 mg/kg IP), A-1331852 (25 mg/kg PO), and the combination of an ADC (3 mg/kg) with A-1331852 (25 mg/kg). Error bars represent standard error of the mean. (B) Immunohistochemical analysis of B7-H3 and p53 at study termination.

To confirm target antigen expression and p53 status in the tumor tissues, samples were collected at day 14 and day 28 of the experiment and B7-H3 and p53 protein was evaluated by immunohistochemistry. We did not appreciate any differences in B7-H3 expression in C4-2B or C4-2B *TP53* KO tumors treated with MGC018 (**Figure 6B**). As expected, p53 staining revealed nuclear p53 expression in C4-2B tumors but absence in C4-2B *TP53* KO tumors (**Figure 6B**). Moderate body weight loss was observed in animals treated with A-1331852, singly or combined with ADC during the first two weeks of the experiment (**Figure S10B**), suggesting that the safety of this drug might be improved by using A-1331852 as an ADC payload. Importantly, the combination of MGC018 with A-1331852 did not accelerate weight loss (**Figure S10B**). Collectively, these findings demonstrate that the addition of a Bcl-xL inhibitor to an ADC bearing a DDD payload can improve tumor responses in both p53-proficient and p53-deficient PC.

## DISCUSSION

The development and translation of effective precision therapies for mCRPC are necessary to alter the course of this highly aggressive disease. Tumor responses to anticancer treatments are often improved by combining cytotoxic agents that operate through diverse mechanisms of action (MoA), and the tolerability of these agents can be enhanced with the use of more effective tumor delivery systems. Antibody drug conjugates (ADCs) offer a unique platform for the development of custom large molecules that target single or multiple tumor-associated antigens and deliver single or multiple cytotoxins (payloads) to the tumor site. Rationally designed combinatorial ADC-based regimens, including simultaneous or sequential ADC co-administration, ADCs combined with systemic agents, and dual-payload ADCs, have the potential to make a clinical impact on the management of mCRPCs.

Most ADC targets are tumor-associated antigens with high abundance in cancer cells and limited normal tissue expression (4–8,34). When ADC affects healthy cells expressing low levels of target antigen, systemic toxicity may arise. Such on-target ADC effects can potentially be reduced by the use of two ADCs targeting different surface antigens, each administered at lower doses, as fewer normal tissues would co-express two tumor-associated targets. In this work we ascertained the co-expression of PC surface antigen(s) in malignant and in benign tissues to strategically select tumor restricted combinations for mCRPC. mCRPC is a phenotypically heterogeneous disease with tumor subtypes marked by distinct transcriptional programs (24,25) and cell surface antigen expression (12,17,35–37). Our analysis of co-expression patterns of PC associated surface antigens across four mCRPC molecular phenotypes demonstrated that B7-H3(*CD276*), PSMA(*FOLH1*), and STEAP1 were frequently co-expressed in the same cells in the tumors with sustained AR signaling. Importantly, co-expression of either explored antigen pair is limited in normal tissues. Thus, our data provides the rationale for the use of ADCs co-targeting B7-H3 and STEAP1, STEAP1 and PSMA, and B7-H3 and PSMA in AR-active tumors which represent about 80% of mCRPCs (24,25).

Purposeful selection of ADC payloads and incorporation of complementary drugs provide another opportunity to improve treatment efficacy. Most ADCs that have been investigated in PC clinical trials employed microtubule disrupting monomethyl auristatin E (MMAE) as a payload, and none of these ADCs demonstrated a satisfactory therapeutic window in mCRPC patients (6). It remains unclear whether MMAE is an optimal cytotoxic agent to control the growth of mCRPCs that were previously exposed to taxane therapy and could have developed resistance mechanisms. Our screening of 23 small molecules from three major ADC payload classes determined that AR-active mCRPC cells had greater average responses to DNA-damaging payloads compared to responses to microtubule targeting drugs. We also found that a NEPC model had remarkable responses to several innovative payloads, including the Bcl-xL inhibitor A-1331852. Further exploration of this result in a larger panel of NEPC models might enable the development for A-1331852 bearing ADCs for this mCRPC subtype that currently has limited treatment options beyond platinum-based chemotherapy.

Through the screening of 78 payload combinations, we identified synergism between DNA-damaging payloads (DDDs) and A-1331852. We find that exposure to this payload combination initiates the intrinsic p53-dependent apoptosis pathway, and that Bcl-xL (*BCL2L1*) expression and p53 activity associate with maximal cytotoxicity. Although synergism was observed in both p53-deficient and p53-proficient PC models, isolated genetic *TP53* knockout reduced cell sensitivity to five DDDs and three DDD bearing ADCs, as well as their combinations with A-1331852. In a prior study assessing the efficacy of a B7-H3 ADC with a pyrrolobenzodiazepine (PBD) payload in 26 treatment-resistant PC PDXs, wildtype *TP53* status predicted non-responsiveness (13). However, *TP53* loss in mCRPC is generally not an isolated event and other genomic and phenotypic features may contribute to ADC efficacy and synergism. Further investigation into p53 status as a biomarker of response to genotoxic ADCs and the influence of other genomic alterations including *BRCA2* and *RB1* mutations are required to direct a precision medicine-based approach to ADC therapy.

We have confirmed that Bcl-xL inhibitor A-1331852 enhances antitumor activity of genotoxic payloads by combining B7-H3-seco-DUBA ADC with unconjugated A-1331852 using *in vivo* C4-2B and C4-2B *TP53 KO* tumor models. These findings have implications for the clinical development of next-generation ADC combinations. Synergism between genotoxic payloads and A-1331852 could be exploited by the development of dual-payload ADCs for the treatment of AR-active and NE mCRPCs. Tumor specificity of these drugs could be further improved by combining two ADCs bearing DDD and A-1331852 payloads. Although concurrent administration of two synergistic ADCs may significantly reduce off-tumor toxicity, it will be crucial to assess clinical pharmacokinetics and tumor penetration of two large molecules in this scenario.

In summary, we have introduced the concept of co-targeting mCRPCs with ADCs bearing synergistic payloads. We have characterized the co-expression of clinically relevant cell surface targets B7-H3, PSMA, and STEAP1 in mCRPC and identified synergism between genotoxic payloads and the Bcl-xL inhibitor payload A-1331852 and others that are undergoing further validation. These findings establish the initial foundation for future clinical development of effective ADC combinations for mCRPC.

## MATERIALS AND METHODS

### Cell lines and Materials

22Rv1 (RRID:CVCL_1045), LNCaP (RRID:CVCL_0395), C4-2B (RRID:CVCL_4784), VCaP (RRID:CVCL_2235), and 293T (RRID:CVCL_0063) cell lines were purchased from the ATCC. Cells were maintained in a 37°C incubator with 5% CO_2_ and grown in medium supplemented with FBS and other additives as recommended by ATCC. LuCaP cell lines were generated by resecting the tumor implant, dissociating cells by enzymatic digestion and plating cells in DMEM medium with 10% FBS or with various additives used in organoid medium (38). MSKCC EF1 (derived from the organoid line MSKCC-CaP4(35)) were maintained in RPMI medium supplemented with 10% FBS, 100 U/mL penicillin and 100 μg/mL streptomycin, and 4 mmol/L GlutaMAX. FreeStyle^TM^ 293-F (RRID:CVCL_D603) cells were purchased from Thermo and maintained in serum free FreeStyle^TM^ 293 Expression Medium on an orbital shaker in 37°C incubator with a humidified atmosphere of 8% CO2. Cell lines were cultured no more than 3 weeks after thawing prior to use in described experiments. Cell lines underwent DNA fingerprint (STR) confirmation and routine mycoplasma testing (R&D Systems CUL001B) via Fred Hutch Research Cell Bank Services. MGC018, vobramitamab duocarmazine, vobra duo) was provided by MacroGenics.

### Immunoblotting

Lysates were prepared in RIPA buffer (Millipore) supplemented with Pierce^TM^ Protein inhibitor cocktail (Thermo), pelleted to remove debris, then quantified by BCA assay (Thermo). Extracts were fractionated by SDS-PAGE and transferred to a PVDF membrane. Membranes were blocked with 5% nonfat milk in TBST (TBS + 1% Tween 20) for 1 hour while shaking, then incubated with primary antibodies at 4°C for 16 hours. Primary antibodies targeting AR (Abcam Cat# ab133273, RRID:AB_11156085, 1:5000), p53 (Santa Cruz Biotechnology Cat# sc-126, RRID:AB_628082, 1:500), Bcl-2 (Santa Cruz Biotechnology Cat# sc-7382, RRID:AB_626736, 1:500), Bcl-xL (Cell Signaling Technology Cat# 2764, RRID:AB_2228008, 1:1000), PARP1 (Thermo Fisher Scientific Cat# 436400, RRID:AB_2532215, 1:1000) cleaved caspase-3 (Cell Signaling Technology Cat# 9661, RRID:AB_2341188, 1:1000), B7-H3 (Sigma SAB5500011, 1:5000), PSMA (Agilent Cat# M3620, RRID:AB_2106450, 1:1000), STEAP1 (Cell Signaling Technology Cat# 88677, RRID:AB_2800128, 1:1000), and GAPDH ((Santa Cruz Biotechnology Cat# sc-47724, RRID:AB_627678, 1:5000) were used. Membanes were washed three times with TBST and incubated with horseradish peroxidase (HRP)-conjugated anti-mouse or anti-rabbit secondary antibody for 1 hour at room temperature. Blots were washed three times with TBST and developed with Immobilon Western Chemiluminescent HRP Substrate (Millipore). Blot images were acquired with a ChemiDoc MP Imaging System (Bio-Rad).

### Payload screening

Twenty-three compounds used in the screen are described in Table S1. Staurosporine (APExBIO A8192) at 1 μM final concentration was used as positive control. All drugs were reconstituted in DMSO and were manually arrayed in a 384-well master plate at concentrations 500 fold greater than final concentration. Drug screening was performed at Fred Hutchinson Genomics and Bioinformatics Shared Resource High Throughput Screening lab. Sub-microliter aliquots of compounds were applied to 384 well culture plates (Corning 3570BC) using a Beckman Coulter Biomek i7 liquid handling instrument fitted with a V&P Scientific 384 Pin Tool applicator. 1-10 × 10^3^ cells in suspension were applied to culture plates containing drug aliquots with a BioTek MicroFlo Select dispense instrument to the final volume of 30ul and incubated for 3 days. To assess cell viability 30 μl of CellTiter Glo 2.0 (Promega) reagent was applied to culture plates with a BioTek MicroFlo Select dispense instrument. Plates were incubated for 20 minutes on a platform shaker and then read using a PerkinElmer EnVision 2104 Multilabel reader.

### Drug dose response curves

Cells were seeded at 5-20 × 10^3^ cells (100 μL) per well in 96-well, tissue culture–treated clear bottom white plates (Corning). Cells were treated with serial dilutions in replicates of 3, at 37°C for 72 hours for free toxin dose response or for 96 hours for ADC dose response. Cell viability was determined using the CellTiter-Glo 2.0 Assay (Promega).

### Generation of B7-H3, PSMA, and STEAP1 knockout cell lines

B7-H3/*CD276*, PSMA/*FOLH1*, and STEAP1 knockout cells were generated by transient transfection of cells with a pool of PX458 (Addgene #48138) plasmids each expressing one sgRNA targeting sequence. B7-H3/*CD276* sgRNAs: (1) 5’-GTGGTCACGTTGCCAGTCAG-3’; (2) 5’-CTGGTGCACAGTTTCACCGA-3’; (3) 5’-CACAGGGCAACGCATCCCTG-3’. PSMA/*FOLH1* sgRNAs: (1) 5’-TTATAGGCGTGGAATTGCAG-3’; (2) 5’-GGAGAGAAAGCACTGAAAGG-3’; (3) 5’-GGTACACAACCTAACAAAAG-3’; (4) 5’-CTGTTGTTCATGAAATTGTG-3’. STEAP1 sgRNAs: (1) 5’-ATAGTCTGTCTTACCCAATG-3’; (2) 5’-CCTTTGTAGCATAAGGACAC-3’; (3) 5’-ATCCACTTATCCAACCAATG-3’; and (4) 5’-CATCAACAAAGTCTTGCCAA-3’. 48–72 hours after transfection, GFP-positive cells were singly sorted on a Sony SH800 Cell Sorter into a 96-well plate and clonally expanded.

### Generation of *TP53* knockout lines

SgRNA (5’-CCCCGGACGATATTGAACAA-3’) was first cloned into the plentiCRISPRv2-blast vector (RRID:Addgene_98293). Lentivirus carrying the construct was then used to transduce C4-2B and LuCaP 189.4 cells, which were subjected to blasticidin selection for 5–7 days. Subsequently, C4-2B cells were plated at a single-cell density to enable clonal expansion. The resulting colonies were expanded and analyzed by western blot. Genomic junctions were characterized by amplifying the sgRNA-targeted regions with sequence-specific primers, followed by Sanger sequencing.

### PSMA and STEAP1 scFv-Fc expression plasmids

gBlocks (IDT) encoding J591 or DSTP3086S scFv (VL-[G4S]3-VH) were cloned into SfiI site of TGEX-SCBlue mammalian expression vector (Antibody Design Labs) using HiFi DNA Assembly (NEB). Resulting TGEX-FOLH1 and TGEX-STEAP1 constructs were analyzed by Sanger sequencing.

### Production of PSMA and STEAP1 scFv-Fc antibodies

Recombinant scFv-Fc antibodies were produced using FreeStyle^TM^ 293 Expression system (Thermo). Briefly, 293-F cells were transfected with TGEX-FOLH1 or TGEX-STEAP1 expression vectors using FreeStyle^TM^ MAX Reagent, and grown in serum-free FreeStyle™ 293 Expression medium for 7 days. Conditioned media was clarified by centrifugation and filter sterilized using Millipore 0.22 μm SteriCup vacuum filtration units. Clarified supernatant was passed through Cytiva HiTrap Protein G HP column, and then bound antibodies were eluted with glycine-HCl buffer (0.1 M glycine, 0.1 M NaCl, pH 2.7), neutralized, and dialyzed against PBS using Slide-A-Lyzer dialysis cassettes (Thermo). After dialysis antibodies were concentrated on 10K MWCO Amicon Ultra-15 centrifugal filter units. Antibody preparations were characterized by SDS-PAGE analysis; protein concentrations were mesured by BCA assay (Thermo).

### Antibody conjugation

Conjugation of PSMA scFv-Fc with Tesirine (*Mal-PEG8-Val-Ala-PABC*-SG3249) or Deruxtecan (*MC-GGFG*-DXd) linker-payload (LP) was performed by NJ Bio, Inc. The antibody; 5 mg/mL in 1X PBS; was reacted by 10-fold molar excess of tris (2-carboxyethyl) phosphine (TCEP). The reaction solution was incubated at 37°C for 90 minutes. Then, 10% v/v Dimethylacetamide (DMA) was added to the solution, followed by 15-fold molar excess of the LP, incubated at RT for 3 hours. The LP was prepared at 10 mM in DMA. To remove the excess LP and aggregation, size exclusion chromatography (SEC) was applied. A Hiload 16/600 Superdex 200 column was equilibrated by 1x PBS buffer, prior to injecting the rection solution. After SEC purification, the ADC was filtered using PES 0.22μm filter and store at 4°C. The DAR was calculated via LC-MS using 0.5 μg of reduced material injected over RP-LC/MS (1000 Å, 8 µm, 2.1 x 50 mm).

Conjugation of STEAP1 scFv-Fc with Deruxtecan LP was performed using Antibody Deruxtecan Conjugation Kit (CellMosaic DCM11431) according to manufacturer protocol. DAR was calculated via HIC HPLC analysis (CellMosaic).

### Flow cytometry

Parental C4-2B and 22Rv1, as well as PSMA KO or STEAP1 KO cell lines were dissociated non-enzymatically with Versene-EDTA into single cell suspensions. 0.5 mln cells were washed trice with monoclonal antibody wash (MW, 1x PBS + 0.1% FBS + 0.1% NaN_3_) and resuspended in 200 μl of 5 μg/ml of PSMA scFv-Fc or STEAP1 scFv-Fc and incubated at 4°C on ice for 1 hour. Cells were then washed with MW, incubated with PE anti-human secondary antibody (BioLegend Cat# 410707, RRID:AB_2565785) at 4°C on ice for 30 minutes, washed with MW, acquired on a SH800 (Sony), and analyzed with FlowJo (RRID:SCR_008520).

### Cell line-derived xenograft studies

Animal care and studies were performed in accordance with an approved Fred Hutchinson Cancer Center Institutional Animal Care and Use Committee protocol and Comparative Medicine regulations. 2 × 10^6^ C4-2B or C4-2B *TP53 KO* cells were suspended in 100 μL of cold 50% Matrigel (Corning)/PBS and implanted by injection subcutaneously into 8-10 weeks old male NSG mice (NOD-SCID-IL2Rγ-null, RRID:BCBC_4142). Animals were assigned to experimental groups using simple randomization, enrolled into experiment when tumors reached 100 mm^3^, and treated by intraperitoneal injection and oral gavage at the frequency and with the doses described. MGC018 was formulated in PBS and administered intraperitoneally (100 μl, at 3 mg/kg) once a week for two weeks. A-1331852 was formulated in 2.5% DMSO, 10% ethanol, 27.5% PEG 300, and 60% Phosal 50 PG. A-1331852 doses (100 μl, at 25 mg/kg) were administered by oral gavage once a day, with five days on-two days off regimen for 2 weeks. Mice were monitored every other day for tumor growth, weight, and body condition score. All enrolled animals finished the study, i.e., there was no attrition.

### Multiplexed Immunofluorescent staining (mIF)

Primary and secondary antibodies used for mIF are listed in Table S2.

Formalin-fixed paraffin-embedded tissues were baked for 1 hour at 65°C. The slides were then dewaxed and stained on a Leica BOND RX stainer (Leica, Buffalo Grove, IL) using Leica Bond reagents for dewaxing (Dewax Solution), antigen retrieval/antibody stripping (Epitope Retrieval Solution 2) and rinsing after each step (Bond Wash Solution). Antigen retrieval and antibody stripping steps were performed at 100°C, with all other steps at ambient temperature. Endogenous peroxidase was blocked with 3% H2O2 for 5 minutes followed by protein blocking with TCT buffer (0.05M Tris, 0.15 M NaCl, 0.25% Casein, 0.1% Tween 20, 0.05% ProClin300 pH 7.6) for 10 minutes. The first primary antibody (Position 1) was applied for 60 minutes, followed by the secondary antibody (1X Opal Anti-Ms + Rb HRP Polymer, Akoya Biosciences) application for 20 minutes, then the application of the tertiary TSA-amplification reagent (Akoya Biosciences OPAL fluor) for 20 minutes. A high-stringency wash was performed after the secondary and tertiary applications using a high-salt TBST solution (0.05M Tris, 0.3M NaCl, and 0.1% Tween-20, pH 7.2-7.6). The primary and secondary antibodies were stripped with retrieval solution for 20 minutes before repeating the process with the second primary antibody (Position 2), starting with a new application of 3% H2O2. The stripping step was not performed after the final position. Slides were removed from the stainer and stained with a 5µg/mL concentration of DAPI (Sigma D8417) for 5 minutes, rinsed, and coverslipped with Prolong Gold Antifade reagent (Invitrogen/Life Technologies P36930). After curing at room temperature, whole slide images were acquired on the Vectra Polaris Quantitative Pathology Imaging System (Akoya Biosciences, Marlborough, MA). The entire tissue was selected for imaging using Phenochart and multispectral image tiles were acquired using the Polaris. Images were spectrally unmixed using Phenoptics inForm software and exported as multi-image TIF files. The TMA slides were visualized with HALO (Indica Labs). H-scores were generated from the mIF data using the CytoNuclear LCv2.0.6 module and HALO software. Individual cells were classified as having negative, weak, moderate, or strong staining and assigned intensity scores. The intensity scores were then multiplied by the percentage of stained cells for a range of 0–300. The triplicate scores were averaged to generate an averaged H-score for each site. To assess total percentage of positive or double-positive cells in the tumor core percentages of cells with weak, moderate, or strong staining were combined.

### Chromogenic Immunohistochemistry (IHC)

IHC was conducted on the VENTANA Discover Ultra (Ventana Medical Systems) automated platform. Sections were cut at a thickness of 4 µm from FFPE PDX tumors and mounted onto positively charged slides. Prior to onboard deparaffination and staining, slides were incubated at 65C for 1 hour to remove excess paraffin. Slides were then loaded onto the machine, incubated in Discovery Wash Buffer (Roche Diagnostics, 950-510) and heat-induced epitope retrieval (HIER) was conducted in Discovery CC1 solution (Roche Diagnostics, 950-224). Primary antibodies (Sigma SAB5500011, 1:500) and P53 (Nolan lab - Stanford Cat# 453M, RRID:AB_2864403, 1:50) were diluted in Ventana Antibody Diluent with Casein (Roche Diagnostics, 760-219) and applied to the sections. The sections probed for P53 were washed and incubated with Rabbit Anti-Mouse IgGs (Abcam Cat# ab46540, RRID:AB_2614925, 1:200). DISCOVERY anti-Rabbit HQ secondary antibody (Roche Cat# 760-4815, RRID:AB_2811171) was applied to all slides, followed by the application of DISCOVERY Anti-HQ HRP (Roche Diagnostics, 760-4820) enzyme conjugate. The antibody-enzyme conjugate complex was visualized with ChromoMap DAB (Roche Diagnostics, 760-159). The sections were counterstained with Hematoxylin II (Roche Diagnostics, 790-2208) and Bluing Reagent (Roche Diagnostics, 760-2037). Slides were scanned at 40X magnification on the Ventana DP 200 slide scanner (Roche Diagnostics).

### Gene expression and genomic analysis

Published data of bulk flash-frozen needle biopsies from the SU2C-IDT/PCF cohort, bulk tumors from the University of Washington (UW) mCRPC cohort, and LuCaP PDX tumors were sequenced and aligned as described previously (17). In brief, sequencing reads were mapped to the hg38 human genome using STAR v2.7.3a (RRID:SCR_004463). PDX data were also aligned to the mm10 mouse genome. All subsequent analyses were performed in R. PDX sequencing reads deriving from mouse were removed using XenofilteR. Gene level abundance was quantitated using GenomicAlignments and transformed to log_2_ TPM. Previously published TP53 mutation status was determined as described by Frank, et al. (39). Mutation status from samples with purity <0.2 were omitted. The UW mCRPC and LuCaP PDX RNAseq data used in this study are available in the Gene Expression Omnibus repository (GEO) Gene Expression Omnibus (GEO) (RRID:SCR_005012) under accession numbers GSE147250 and GSE199596. SU2C-IDT/PCF RNAseq data are available in the cBioPortal (prad_su2c_2019; https://github.com/cBioPortal/datahub/tree/master/public/prad_su2c_2019.)

### Statistical methods

For each target antigen PSMA, STEAP1, and B7-H3, median differences in average H-scores between pairs of phenotypes were evaluated using Wilcoxon-Mann-Whitney rank sum tests using Holm’s method to adjust for multiple comparisons using the wilcox_test function from the rstatix package (RRID: SCR_021240) in the R programming language (RRID: SCR_001905). This method was also used to evaluate median differences in relative viability between payload classes within each cell line. Intra-patient and intra-tumoral heterogeneity indices were estimated via bootstrapping from the same patient or tumor and evaluating whether pairs of samples had discordant positivity (i.e., one sample <20% and one sample ≥20%). Longitudinal tumor volume measurements (on the logarithmic scale) were analyzed by fitting a linear model with random intercepts for independent animals and fixed effects for day, treatment group, and their interaction using the lmer function from the lmerTest package (RRID: SCR_015656); two-sided p-values for different slopes were used to determine effects of treatment on average daily tumor growth rates.

## Authors’ Contributions

GS - conception and design of the study, acquisition, analysis and interpretation of data, drafting and revision of the manuscript; SF – conceptualization, data interpretation, manuscript revision; RD – data acquisition, WH – models development, manuscript drafting; IC – data analysis, manuscript drafting; RG – data analysis, manuscript drafting and revision; CM – data acquisition, resources supply, manuscript revision; MCH – analysis and interpretation of the data, manuscript revision; PSN – design of the research, data analysis, models supply, manuscript revision; JKL – conceptualization and design, resources, supervision, manuscript drafting and revision. All authors read and approved the final manuscript.

## Acknowledgements

We thank members of the Lee and Nelson research groups for providing advice and input. We thank Phil Corrin for assistance with experiments involving drug screens. We thank MacroGenics, Inc for providing MGC018, vobramitamab duocarmazine. We thank the patients and their families, Evan Yu, Heather Cheng, Bruce Montgomery, Jessica Hawley, Mike Schweizer, Celestia Higano, Andrew Hsieh, Daniel Lin, Funda Vakar-Lopez, Xiaotun Zhang, Martine Roudier, Lawrence True, Meagan Chambers, Eva Corey and the rapid autopsy teams for their contributions to the University of Washington Medical Center Prostate Cancer Donor Rapid Autopsy Program. This research was supported by the Experimental Histopathology Shared Resource, RRID:SCR_022612, of the Fred Hutch/University of Washington/Seattle Children’s Cancer Consortium (P30CA015704). This work was also supported in part by awards from the National Cancer Institute (P50CA097186, R50CA221836; R01CA266452 and R50CA274336), CDMRP awards PC230582 and PC230420, The Department of Defense Prostate Cancer Research Program (W81XWH-14-2-0183), the Prostate Cancer Foundation and the Institute for Prostate Cancer Research.

