## Supplementary material for "Genotoxic antibody-drug conjugates combined with Bcl-xL inhibitors enhance therapeutic efficacy in metastatic castration-resistant prostate cancer": Tables S1, S2, and S5

Table S1. Cytotoxic agents used in the drug screen.

| **Compound name** | **Vendor** | **Catalog #** | **Concentration (high) in uM** | **Concentration (low) in uM** |
| --- | --- | --- | --- | --- |
| 10-Deacetyl-7-xylosyl paclitaxel | MedChemExpress | HY-20584 | 5 | 0.5 |
| PF-06380101 | MedChemExpress | HY-12522 | 0.001 | 0.0001 |
| Mertansine | MedChemExpress | HY-19792 | 0.001 | 0.0001 |
| DM4 | MedChemExpress | HY-12454 | 0.005 | 0.0005 |
| Ispinesib | MedChemExpress | HY-50759 | 0.005 | 0.0005 |
| SG3199 | MedChemExpress | HY-101161 | 0.001 | 0.0001 |
| SJG-136 | MedChemExpress | HY-14573 | 0.005 | 0.0005 |
| PNU-159682 | MedChemExpress | HY-16700 | 0.001 | 0.0001 |
| Daunorubicin (hydrochloride) | MedChemExpress | HY-13062 | 1 | 0.1 |
| DXd | MedChemExpress | HY-13631D | 1 | 0.1 |
| Calicheamicin | MedChemExpress | HY-19609 | 0.001 | 0.0001 |
| Thailanstatin A | MedChemExpress | HY-129589 | 1 | 0.1 |
| Duocarmycin TM | MedChemExpress | HY-107769 | 0.001 | 0.0001 |
| Ansamitocin P-3 | Sigma-Aldrich | A2836 | 5 | 0.5 |
| Camptothecin | Sigma-Aldrich | C9911 | 1 | 0.1 |
| Dolastatin 10 | Sigma-Aldrich | CS-1825 | 1 | 0.1 |
| A-1331852 | Cayman Chemical | 22963 | 5 | 0.5 |
| α-Amanitin | Cayman Chemical | 17898 | 0.5 | 0.05 |
| β-Amanitin | Cayman Chemical | 18142 | 0.5 | 0.05 |
| FK-866 | Cayman Chemical | 13287 | 0.5 | 0.05 |
| CHS-828 | Cayman Chemical | 11021 | 0.05 | 0.005 |
| SN-38 | Selleckchem | S4908 | 0.1 | 0.01 |
| MMAE | Selleckchem | S7721 | 0.001 | 0.0001 |

Table S2. List of primary and secondary antibodies used for multiplexed immunofluorescent staining.

| **Protein target** | **Antibody** | **Host/clone** | **Manufacturer/ Cat#** | **Concentration/ Dilution** | **Secondary/ Cat#** | **Opal Dye/ Cat#** |
| --- | --- | --- | --- | --- | --- | --- |
| 1 | **STEAP1** | Rabbit polyclonal | LS Bio LS-C291740 | 2µg/ml 1:500 | 1X Opal Anti-Ms + Rb HRP ARH1001EA | 570 FP1488001KT |
| 2 | **PSMA** | Mouse 3E6 | Agilent M3620 | 0.157µg/ml 1:1000 | 1X Opal Anti-Ms + Rb HRP ARH1001EA | 690 FP1497001KT |
| 3 | **B7-H3** | Rabbit  EPR20115 | Abcam ab219648 | 0.038 µg/ml 1:15,000 | 1X Opal Anti-Ms + Rb HRP ARH1001EA | 520 FP1487001KT |

Table S5. Components of genotoxic ADCs targeting B7-H3, PSMA, and STEAP1.

| **ADC Name** | **ADC**  **Target** | **Antibody**  **Format** | **ADC Payload** | **ADC Linker** | **DAR** |
| --- | --- | --- | --- | --- | --- |
| B7-H3-seco-DUBA  (MGC018)  (vobramitamab duocarmazine)  (vobra duo) | **B7-H3** | IgG1 | Seco-DUBA | Val-Cit | 2.7 |
| PSMA-SG3249 | **PSMA** | scFv-Fc | SG3249  (Tesirine) | Mal-PEG8-Val-Ala-PABC | 3 |
| STEAP1-DXd | **STEAP1** | scFv-Fc | DXd | MC-GGFG | 1.09 |
