## Supplementary material for "Genotoxic antibody-drug conjugates combined with Bcl-xL inhibitors enhance therapeutic efficacy in metastatic castration-resistant prostate cancer": Figures S1-S10 and Legends

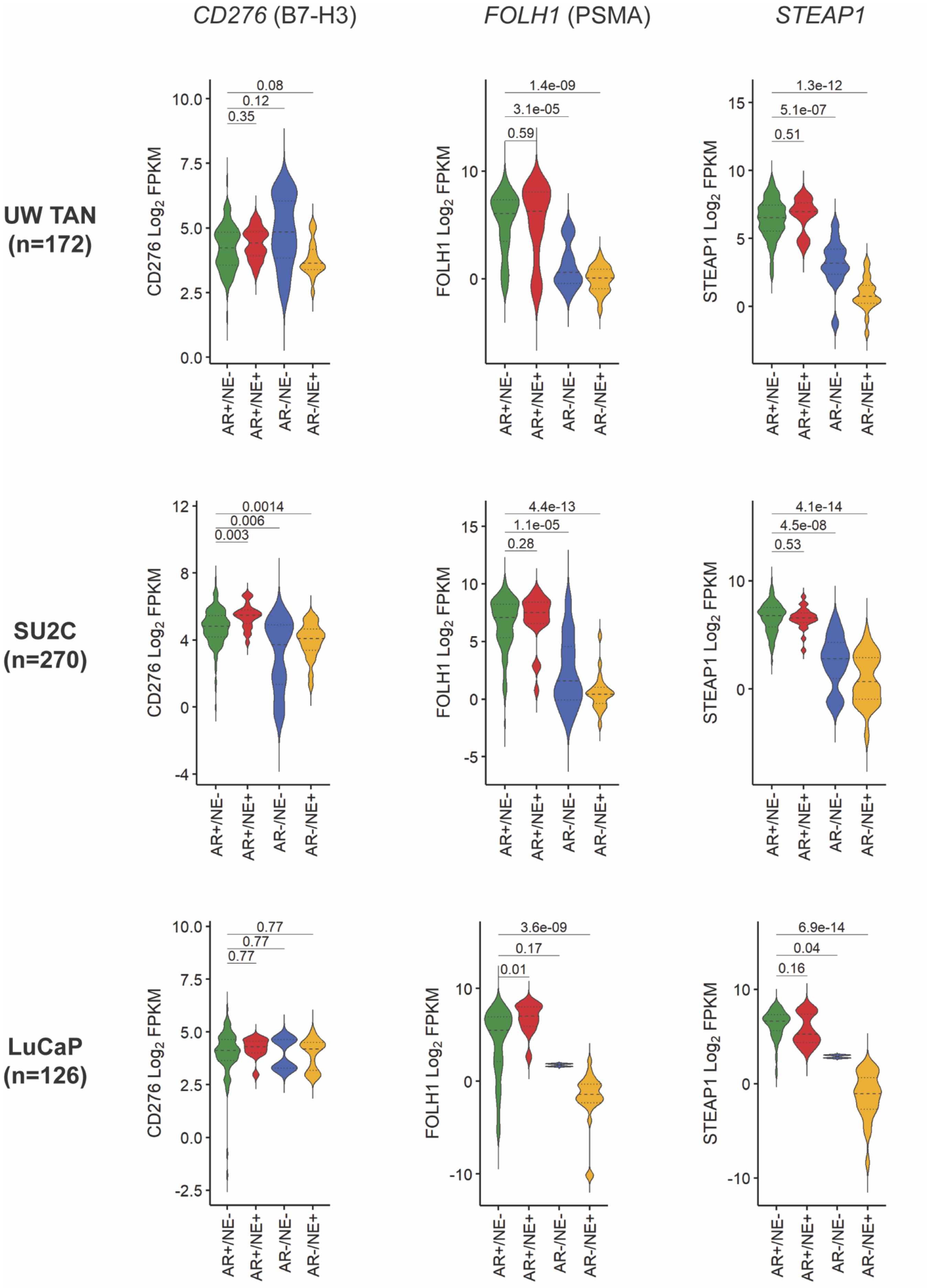
CD276 (B7-H3)
FOLH1 (PSMA)
STEAP1
UW TAN (n=172)
SU2C (n=270)
LuCaP (n=126)
CD276 Log2 FPKM
FOLH1 Log2 FPKM
STEAP1 Log2 FPKM
AR+/NE-
AR+/NE+
AR-/NE-
AR-/NE+
0.08
0.12
0.35
1.4e-09
3.1e-05
0.59
1.3e-12
5.1e-07
0.51
0.0014
0.006
0.003
4.4e-13
1.1e-05
0.28
4.1e-14
4.5e-08
0.53
0.77
0.77
0.77
3.6e-09
0.17
0.01
6.9e-14
0.04
0.16

**Figure S1. *CD276* (B7-H3), *FOLH1* (PSMA), and *STEAP1* expression across mCRPC molecular subtypes.**

Violin plots show *CD276*, *FOLH1*, and *STEAP1* transcript levels in AR+/NE-(green), AR+/NE+(red), AR-/NE-(blue), and AR-/NE+(yellow) tumors from UW TAN, SU2C, and LuCaP cohorts. Results are expressed as log2 fragments per kilobase of transcript per million mapped reads (FPKM). The groups were compared using two-sided Wilcoxon rank tests with Benjamini-Hochberg multiple-testing correction.

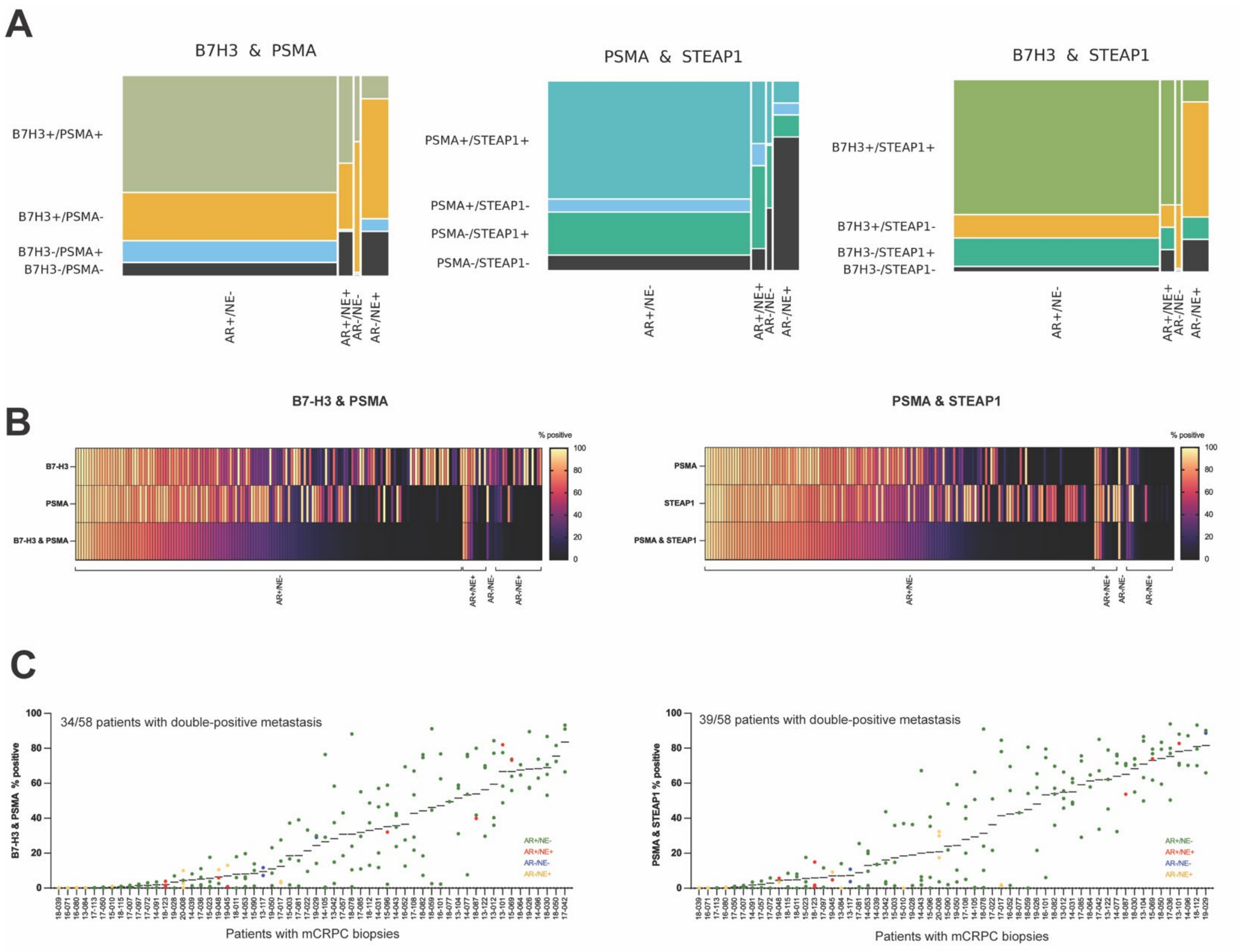


**Figure S2. Co-expression of B7-H3 and PSMA / PSMA and STEAP1 / B7-H3 and STEAP1 in mCRPC tumors and patients.**

(A) Mosaic plot showing mCRPC subtypes scaled to their relative proportions versus marker status pairs scaled to their relative proportions within each subtype. Double positivity is defined as ≥20% positive cells. (B) Heatmaps showing percents of cells staining positively for B7-H3 and PSMA (left) or PSMA and STEAP1 (right) in each individual mCRPC tumor (columns, n=176). (B). Distribution of B7-H3 and PSMA (left) or PSMA and STEAP1 (right) double-positive cells in 176 metastatic tumors within and between 58 patients from UW TAN cohort. Each dot represents a tumor sample; the color codes indicate the molecular subtype – AR+/NE- (green), AR+/NE+ (red), AR-/NE- (blue), and AR-/NE+ (yellow).

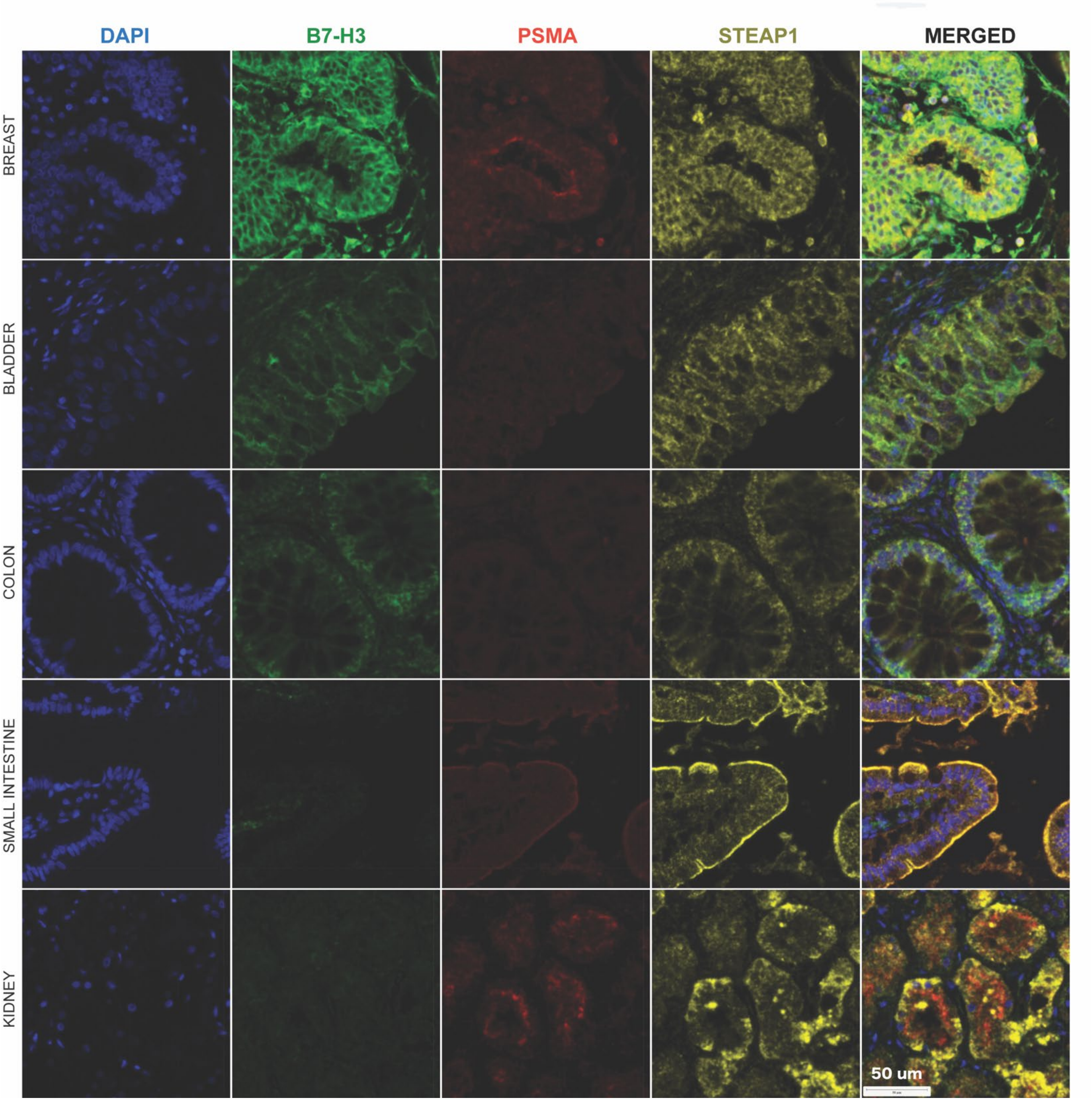


**Figure S3. Normal tissues co-expressing B7-H3, PSMA, and STEAP1.**

Representative TMA images of human breast, bladder, kidney, small and large intestine tissues (FDA999 L206) with membranous B7-H3, PSMA, STEAP1, and nuclear DAPI staining.

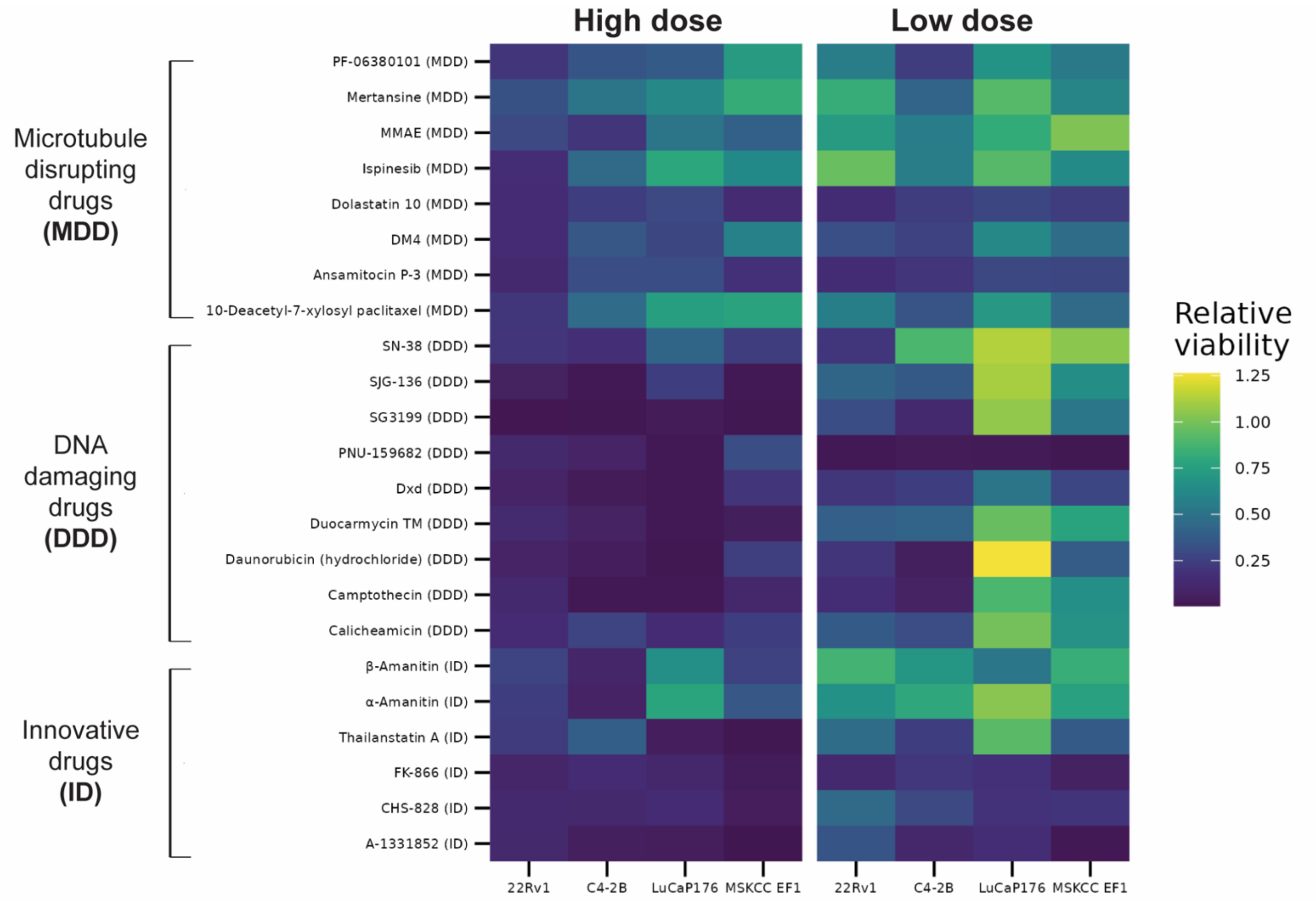


| Line | Group 1 | Group 2 | n 1 | n 2 | Low dose Estimate (95% CI) | Low dose P-value | High dose Estimate (95% CI) | High dose P-value |
|---|---|---|---|---|---|---|---|---|
| **C4-2B** | MDD | DDD | 8 | 9 | 0.09 (0.03, 0.17) | 0.005 | 0.13 (-0.10, 0.28) | >0.9 |
| | MDD | ID | 8 | 6 | 0.03 (-0.08, 0.10) | 0.5 | 0.01 (-0.44, 0.26) | >0.9 |
| | DDD | ID | 9 | 6 | -0.07 (-0.14, 0.00) | 0.18 | -0.10 (-0.46, 0.15) | >0.9 |
| **22Rv1** | MDD | DDD | 8 | 9 | 0.28 (0.17, 0.41) | <0.001 | 0.31 (-0.05, 0.55) | 0.2 |
| | MDD | ID | 8 | 6 | 0.23 (0.09, 0.36) | 0.02 | 0.07 (-0.31, 0.43) | 0.8 |
| | DDD | ID | 9 | 6 | -0.06 (-0.13, 0.03) | 0.11 | -0.24 (-0.47, 0.03) | 0.2 |
| **LuCaP176** | MDD | DDD | 8 | 9 | 0.36 (0.26, 0.60) | 0.003 | -0.27 (-0.58, 0.02) | 0.18 |
| | MDD | ID | 8 | 6 | 0.24 (-0.27, 0.57) | 0.18 | 0.13 (-0.24, 0.63) | 0.5 |
| | DDD | ID | 9 | 6 | -0.10 (-0.64, 0.03) | 0.18 | 0.41 (-0.05, 0.88) | 0.2 |
| **MSKCC EF1** | MDD | DDD | 8 | 9 | 0.41 (0.13, 0.59) | 0.02 | -0.06 (-0.33, 0.26) | >0.9 |
| | MDD | ID | 8 | 6 | 0.44 (0.14, 0.72) | 0.02 | 0.22 (-0.30, 0.52) | >0.9 |
| | DDD | ID | 9 | 6 | 0.02 (-0.12, 0.21) | 0.6 | 0.23 (-0.19, 0.60) | >0.9 |

**Figure S4. Prostate cancer cell lines demonstrate greater response to DNA-damaging drugs (DDD) compared to microtubule-disrupting drugs (MDD).**

Heatmap (top) visualizes relative viability by drug and cell line for high and low doses. RLU – relative luminescence units (relative cell viability). Pairwise comparisons of relative cell viability between payload groups (bottom). The groups were compared using Wilcoxon-Mann-Whitney test.

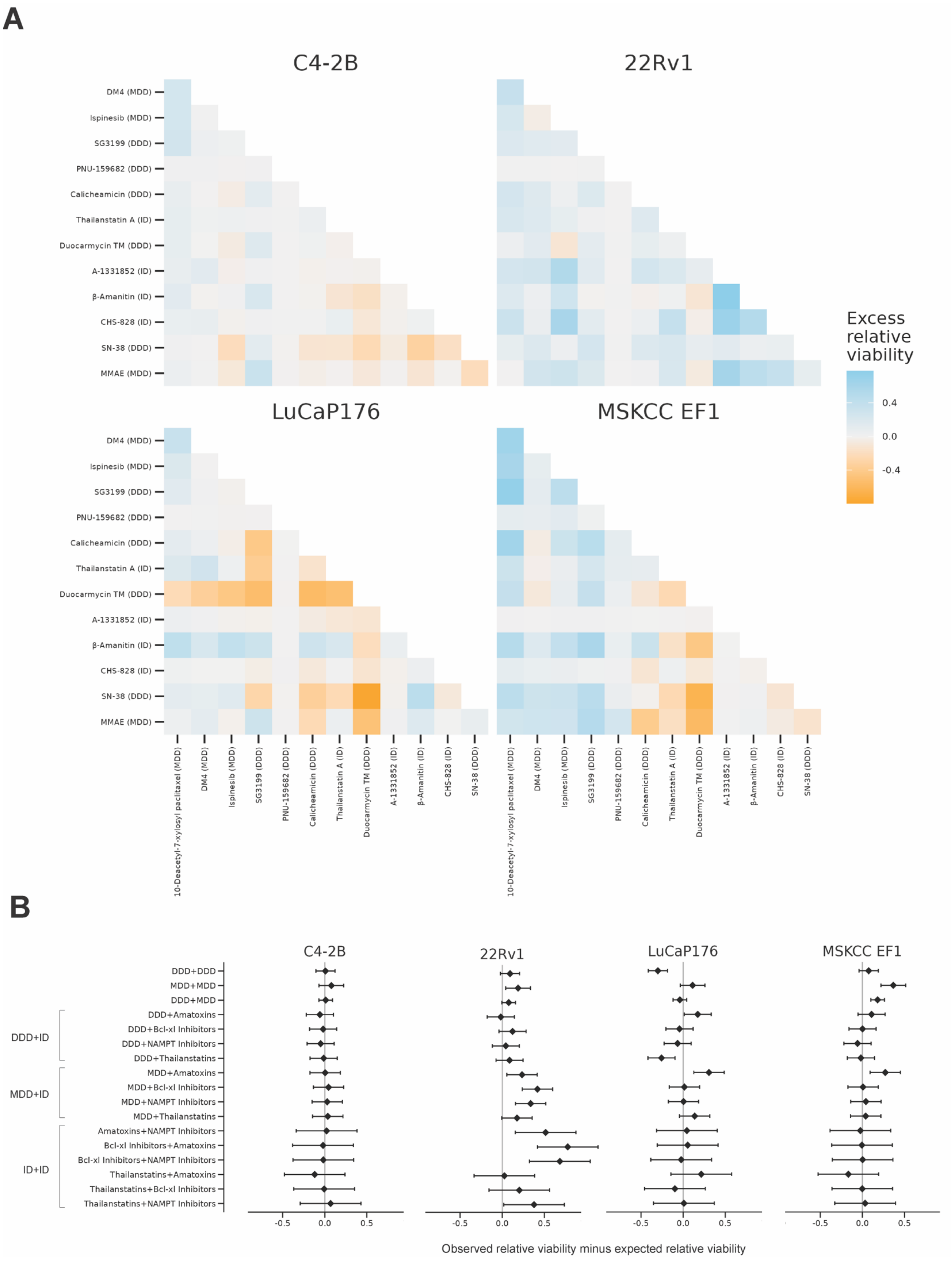
A
C4-2B
22Rv1
LuCaP176
MSKCC EF1
DM4 (MDD)
Ispinesib (MDD)
SG3199 (DDD)
PNU-159682 (DDD)
Calicheamicin (DDD)
Thailanstatin A (ID)
Duocarmycin TM (DDD)
A-1331852 (ID)
β-Amanitin (ID)
CHS-828 (ID)
SN-38 (DDD)
MMAE (MDD)
10-Deacetyl-7-xylosyl paclitaxel (MDD)
Excess relative viability
0.4
0.0
-0.4
B
DDD+DDD
MDD+MDD
DDD+MDD
DDD+Amatoxins
DDD+Bcl-xl Inhibitors
DDD+NAMPT Inhibitors
DDD+Thailanstatins
MDD+Amatoxins
MDD+Bcl-xl Inhibitors
MDD+NAMPT Inhibitors
MDD+Thailanstatins
Amatoxins+NAMPT Inhibitors
Bcl-xl Inhibitors+Amatoxins
Bcl-xl Inhibitors+NAMPT Inhibitors
Thailanstatins+Amatoxins
Thailanstatins+Bcl-xl Inhibitors
Thailanstatins+NAMPT Inhibitors
DDD+ID
MDD+ID
ID+ID
-0.5
0.0
0.5
Observed relative viability minus expected relative viability

**Figure S5. Combination payload screening prioritizes candidates for synergy assessment.**

(A) Heatmaps showing excess relative viability of C4-2B, 22Rv1, LuCaP 176, and MSKCC EF1 cells exposed to single payloads at low dose and payload combinations. Excess viability was calculated as *observed viability* ($viability_{drug1+drug2}$) - *expected viability* ($viability_{drug1}$ * $viability_{drug2}$). (B). Excess relative viability means for the combinations between payload groups and classes in four cell lines.

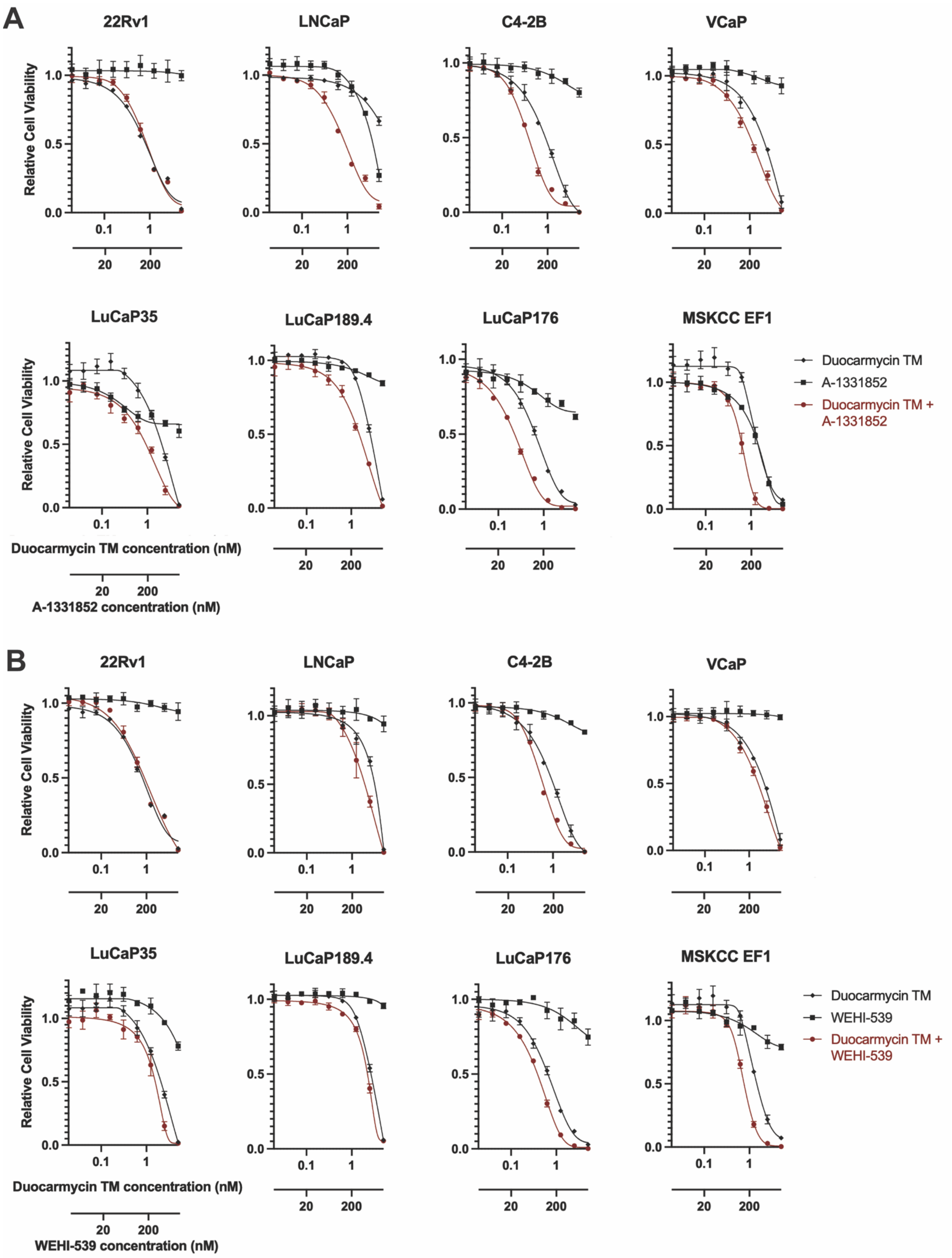
A
22Rv1
LNCaP
C4-2B
VCaP
LuCaP35
LuCaP189.4
LuCaP176
MSKCC EF1
Relative Cell Viability
0.0
0.5
1.0
0.1
1
20
200
Duocarmycin TM concentration (nM)
A-1331852 concentration (nM)
Duocarmycin TM
A-1331852
Duocarmycin TM + A-1331852
B
22Rv1
LNCaP
C4-2B
VCaP
LuCaP35
LuCaP189.4
LuCaP176
MSKCC EF1
Relative Cell Viability
Duocarmycin TM concentration (nM)
WEHI-539 concentration (nM)
Duocarmycin TM
WEHI-539
Duocarmycin TM + WEHI-539

**Figure S6. Duocarmycin TM and Bcl-xL inhibitors (A-1331852 and WEHI-539) combinations exhibit synergistic cytotoxicity in a panel of prostate cancer cell lines.**

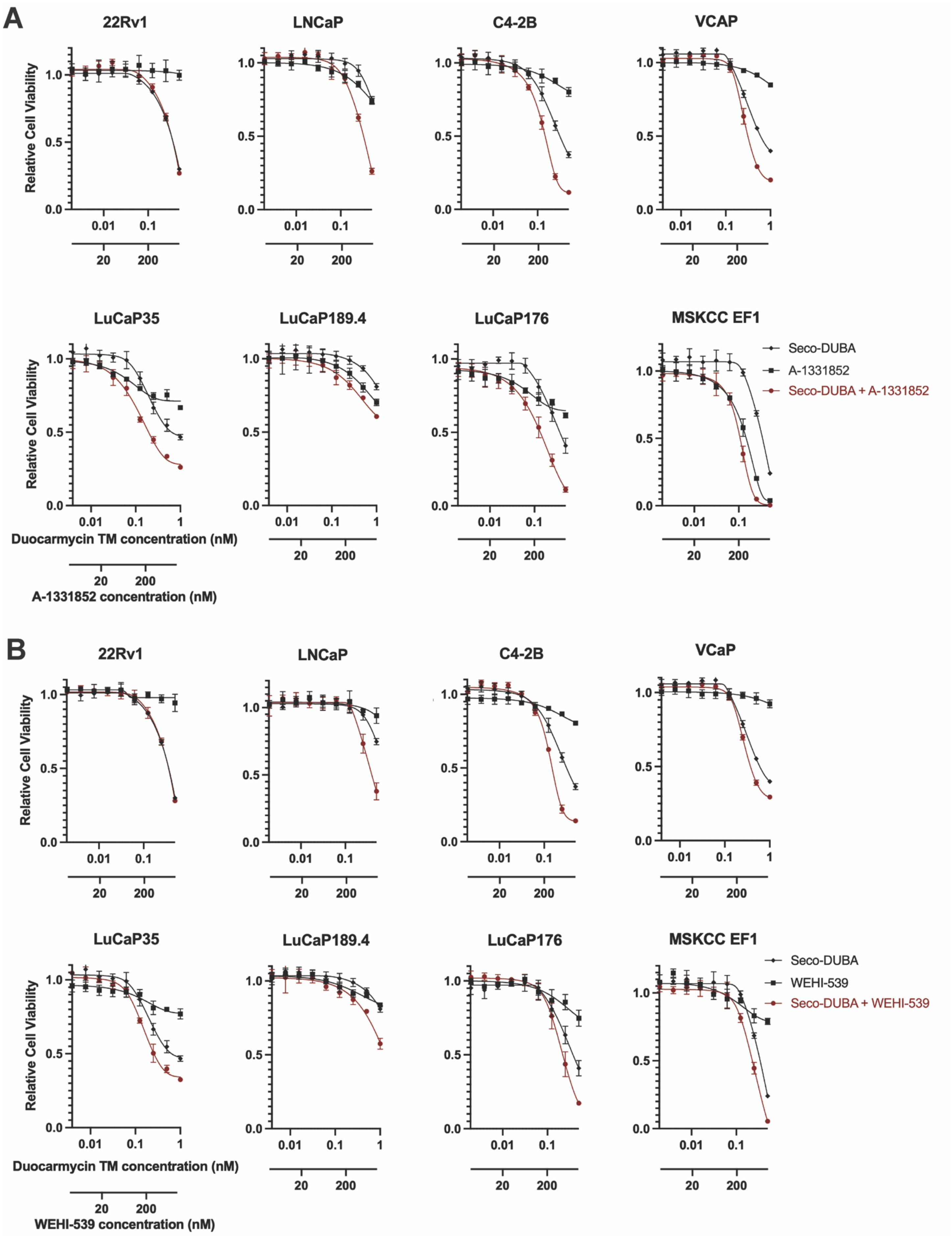
A
22Rv1
LNCaP
C4-2B
VCAP
LuCaP35
LuCaP189.4
LuCaP176
MSKCC EF1
Relative Cell Viability
0.0
0.5
1.0
0.01
0.1
1
20
200
Duocarmycin TM concentration (nM)
A-1331852 concentration (nM)
Seco-DUBA
A-1331852
Seco-DUBA + A-1331852
B
22Rv1
LNCaP
C4-2B
VCaP
LuCaP35
LuCaP189.4
LuCaP176
MSKCC EF1
Relative Cell Viability
Duocarmycin TM concentration (nM)
WEHI-539 concentration (nM)
Seco-DUBA
WEHI-539
Seco-DUBA + WEHI-539

**Figure S7. Seco-DUBA and BCL-XL inhibitors (A-1331852 and WEHI-539) combinations exhibit synergistic cytotoxicity in a panel of prostate cancer cell lines.**

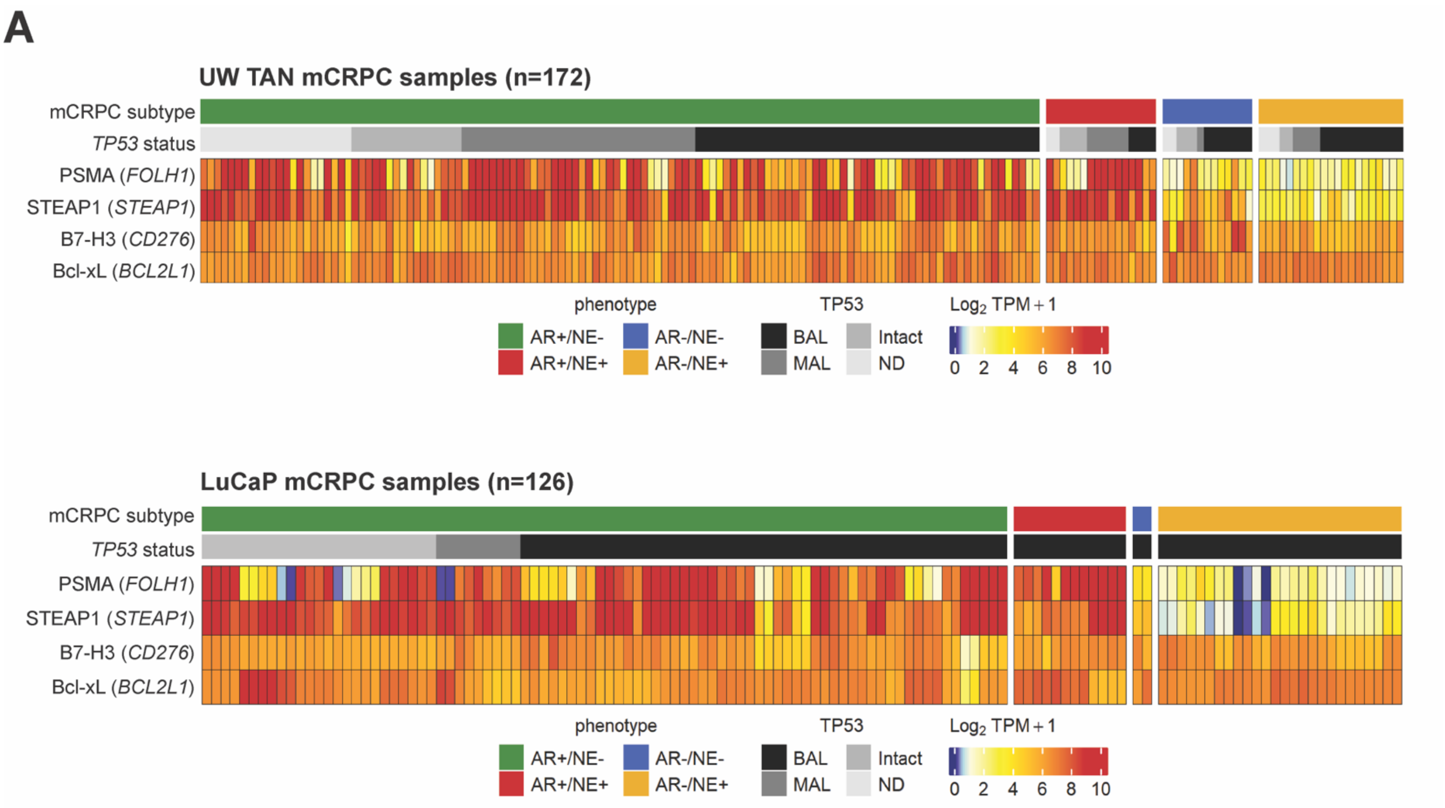


**B**

| Dataset | BAL | Intact | MAL | ND | Total | %BAL |
|---|---|---|---|---|---|---|
| SU2C | 95 | 68 | 91 | 16 | 270 | 35% |
| AR+/NE- | 71 | 58 | 69 | 12 | 210 | 34% |
| AR+/NE+ | 10 | 3 | 10 | 0 | 23 | 43% |
| AR-/NE- | 3 | 4 | 5 | 2 | 14 | 21% |
| AR-/NE+ | 11 | 3 | 7 | 2 | 23 | 48% |
| UW TAN | 73 | 25 | 45 | 29 | 172 | 42% |
| AR+/NE- | 50 | 16 | 34 | 22 | 122 | 41% |
| AR+/NE+ | 4 | 4 | 6 | 2 | 16 | 25% |
| AR-/NE- | 7 | 3 | 1 | 2 | 13 | 54% |
| AR-/NE+ | 12 | 2 | 4 | 3 | 21 | 57% |
| LuCaP | 92 | 25 | 9 | 0 | 126 | 73% |
| AR+/NE- | 52 | 25 | 9 | 0 | 86 | 60% |
| AR+/NE+ | 12 | 0 | 0 | 0 | 12 | 100% |
| AR-/NE- | 2 | 0 | 0 | 0 | 2 | 100% |
| AR-/NE+ | 26 | 0 | 0 | 0 | 26 | 100% |

**Figure S8. *TP53* genomic alterations in mCRPCs expressing B7-H3 (*CD276*), PSMA (*FOLH1*), STEAP1, and Bcl-xL (*BCL2L1*).**

(A) Heatmap showing *FOLH1, STEAP1, CD276,* and *BCL2L1* transcript abundance, as well as *TP53* genomic status in UW TAN and LuCaP mCRPC specimens. Transcript levels are shown as $Log_2$ TPM + 1. BAL – biallelic loss, MAL – monoallelic loss, ND – no data. (B) Fractions of tumors with and without

TP53 genomic alterations in SU2C, UW TAN, and LuCaP cohorts across 4 mCRPC molecular subtypes.

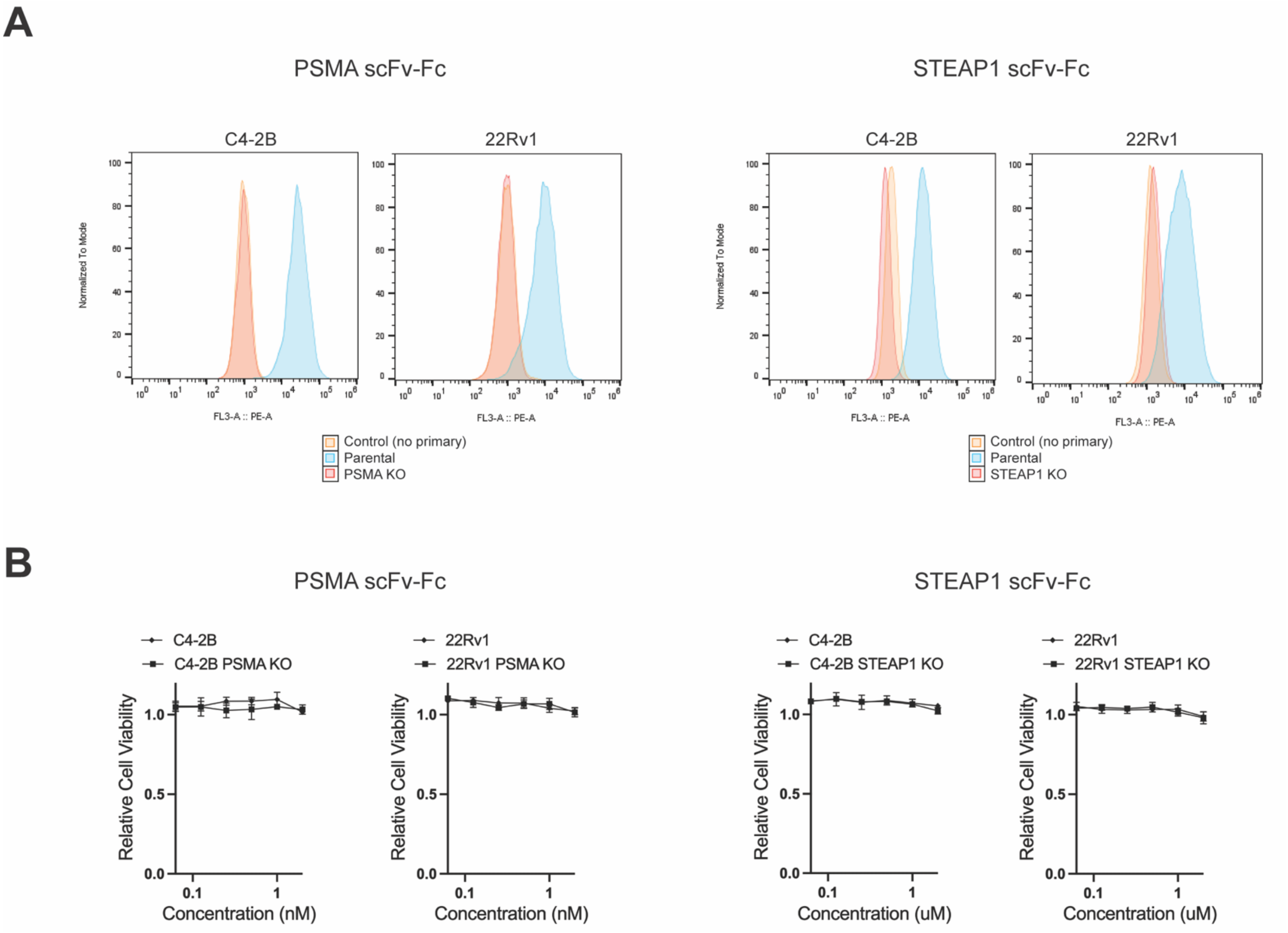


**Figure S9. Quality control of PSMA and STEAP1 scFv-Fc antibodies.**

(A) PSMA scFv-Fc or STEAP1 scFv-Fc binding to the target antigens determined by flow cytometry in parental and antigen KO C4-2B and 22Rv1 cells. (B) Viability of C4-2B and 22Rv1 cells exposed to naked PSMA scFv-Fc or STEAP1 scFv-Fc.

**A**

| **Group * Day** | C4-2B exp(Beta) | C4-2B 95% CI[1] | C4-2B p-value | C4-2B *TP53 KO* exp(Beta) | C4-2B *TP53 KO* 95% CI[1] | C4-2B *TP53 KO* p-value |
|---|---|---|---|---|---|---|
| B7-H3 – seco-DUBA* Day | 0.95 | 0.94, 0.96 | <0.001 | 0.97 | 0.96, 0.98 | <0.001 |
| A1331852 * Day | 0.97 | 0.96, 0.98 | <0.001 | 0.99 | 0.98, 1.00 | 0.065 |
| B7-H3 – seco-DUBA+A1331852 * Day | 0.94 | 0.93, 0.94 | <0.001 | 0.96 | 0.95, 0.96 | <0.001 |

**B**

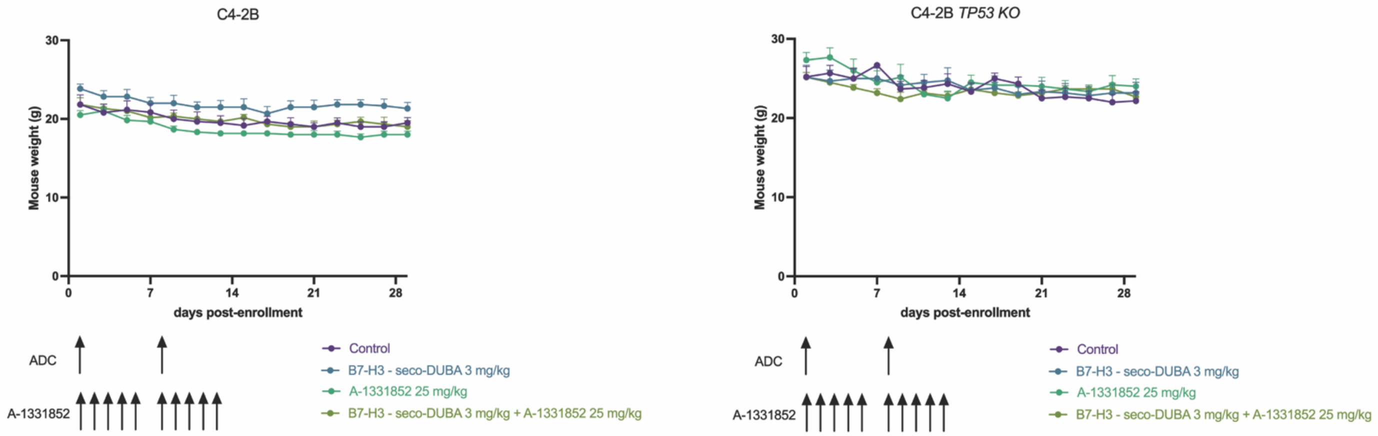


**Figure S10. Analysis of volumetric changes in CDX tumors and body weight changes in animals from control and treatment C4-2B and C4-2B *TP53 KO* groups.**

(A) Statistical analysis for the estimated tumor volume growth rate with tests relative to the control. CI[1] – confidence interval. (B) Average body weights per group throughout the experiment. Error bars represent standard error of the mean.
